# Impact of inoculation practices on microbiota assembly and community stability in a fabricated ecosystem

**DOI:** 10.1101/2023.06.13.544848

**Authors:** Hsiao-Han Lin, Marta Torres, Catharine A. Adams, Peter F. Andeer, Trenton K. Owens, Kateryna Zhalnina, Lauren K. Jabusch, Hans K. Carlson, Jennifer V. Kuehl, Adam M. Deutschbauer, Trent R. Northen, N. Louise Glass, Jenny C. Mortimer

**Author notes:** Corresponding author: Jenny C. Mortimer.

## Abstract

Studying plant-microbe-soil interactions is challenging due to their high complexity and variability in natural ecosystems. While fabricated ecosystems provide opportunities to recapitulate aspects of these systems in reduced complexity and controlled environments, inoculation can be a significant source of variation. To tackle this, we evaluated how different bacteria inoculation practices and plant harvesting time points affect the reproducibility of a microbial synthetic community (SynCom) in association with the model grass *Brachypodium distachyon*. We tested three microbial inoculation practices: seed inoculation, transplant inoculation, and seedling inoculation; and two harvesting points: early (14-day-old plants) and late (21 days post-inoculation). We grew our plants and bacterial strains in sterile devices (EcoFABs) and characterized the microbial community from root, rhizosphere, and sand using 16S ribosomal RNA gene sequencing. The results showed that inoculation practices significantly affected the rhizosphere microbial community only when harvesting at an early time point but not at the late stage. As the SynCom showed a persistent association with *B. distachyon* at 21 days post-inoculation regardless of inoculation practices, we assessed the reproducibility of each inoculation method and found that transplant inoculation showed the highest reproducibility. Moreover, plant biomass was not adversely affected by transplant inoculation treatment. We concluded that bacteria inoculation while transplanting coupled with a later harvesting time point gives the most reproducible microbial community in the EcoFAB-*B. distachyon*-SynCom fabricated ecosystem and recommend this method as a standardized protocol for use with fabricated ecosystem experimental systems.

## INTRODUCTION

Increasing pressure on agricultural systems and environmental degradation have led to examining new methods for developing sustainable agricultural practices in order to achieve global food security goals (United Nation 2022). Much effort is invested nowadays on developing microbial inoculants to replace or supplement fertilizers to improve crop productivity and environmental sustainability (Adesemoye and Kloepper 2009; Tyagi et al. 2022). The vast majority of conventional biological management strategies and scientific studies assessing the impact of biotic/abiotic stresses on biocontrol agents and/or native microbial communities use single microbial agents as inoculants (Khatoon et al. 2020; Oleńska et al. 2020). However, the activity of individual microbial species might differ when the strains are applied as part of a community or consortium (Finkel et al. 2020; Matos and Garland 2005). This is the reason why in recent years the use of synthetic microbial communities (SynComs) has been explored (Liu et al. 2019; McCarty and Ledesma-Amaro 2019; de Souza et al. 2020). SynComs represent systems of reduced complexity that involve co-culturing multiple taxa under well-defined conditions to mimic the structure and function of a microbiome.

From an applied and biotechnological point of view, the SynCom approach has become popular because it can be used to assess how to develop versatile and productive multi-bacterial inoculants for the agriculture sector (McCarty and Ledesma-Amaro 2019). From an ecological point of view, they are a useful tool for a more realistic understanding of the outcomes of multiple biotic interactions where microbes, plants, and the environment are players in time and space of a multidimensional and complex system (Liu et al. 2019; Pradhan et al. 2022; de Souza et al. 2020). SynComs provide functional and mechanistic insights into how plants regulate their microbiomes, and they can be used to assess the impact of biotic and abiotic stresses in microbial communities (McCarty and Ledesma-Amaro 2019; Pradhan et al. 2022; de Souza et al. 2020). Additionally, they are useful for understanding the dynamic interactions within plant-bacteria ecosystems, which is applicable for tool development and identification of candidates for targeted microbiome manipulation (Shayanthan et al. 2022; de Souza et al. 2020).

A successful SynCom inoculant and inoculation methodology needs to address three criteria: persistent plant association, formation of a stable microbial community, and consistency in producing favorable plant phenotypes (Bhardwaj et al. 2014; Mitter et al. 2021). Soil-plant-microbe experimental systems are complex, and bridging the lab-field gap in a replicable and reproducible manner is laborious (Parnell et al. 2016; de Souza et al. 2020). A major challenge in identifying appropriate SynCom inoculants is the scarcity of comparable datasets. The lack of comparable datasets is, in part, due to the absence of standardized systems and protocols (Meisner et al. 2022; York et al. 2022). In an effort to address consistency and to achieve reproducibility, especially for plant traits, researchers have developed different types of fabricated ecosystems (e.g. EcoFAB, Rhizobox, and Rhizotron) (Busch et al. 2006; Ke et al. 2021; Oburger et al. 2013; Sasse et al. 2019; Yee et al. 2021). One of these ecosystems, EcoFABs, has enabled reproducible *Brachypodium distachyon* plant phenotypes in the absence of bacterial inoculant in a multi-laboratory experiment across four different labs (Sasse et al. 2019). Experiments involving bacterial SynComs add an extra layer of complexity to these assays with fabricated ecosystems. Ensuring microbial colonization and predictable long-term plant phenotype are challenges to overcome in these experiments.

SynCom assays need to be accompanied by standardized methods that include SynCom preparation and inoculation procedures, plant stage at which the SynCom is applied to plants, and data analysis pipelines. Comparable data on the impact of plant age, inoculation method and sample harvesting time on microbial community composition is currently scarce. In the future, multi-laboratory experiments using standardized methods to measure reproducibility between fabricated ecosystems inoculated with the same SynCom could provide very valuable information to the field.

Few studies have investigated the effects of inoculation methods on the dynamics of plant-microbe interactions. There are several methods to apply a SynCom to the plant: seed inoculation, transplant inoculation, and seedling root inoculation (Bashan 1998; Kumar et al. 2022; Lopes et al. 2021). The seed inoculation method involves applying a bacterial culture (liquid or powdered) directly to the seeds, and it is widely used in agriculture (O’Callaghan 2016; Rocha et al. 2019; Sarkar et al. 2021). The inoculation upon transplanting method involves microbial inoculation when transferring seedlings from the nursery plate to their permanent growth habitat (e.g. pot, field), and it is commonly used for vegetable growth and lab studies (Herrera Paredes et al. 2018; Hu et al. 2020; Schillaci et al. 2020). The seedling root inoculation involves bacteria inoculation to the root after the plants have been growing for some time in their final habitat (dos Santos et al. 2019; Valetti et al. 2018). How these different inoculation methods affect the formation of a stable microbial community, persistent plant association, and consistency of the plant-microbe ecosystem is poorly understood.

With the aim to answer these questions we conducted a study using fabricated ecosystems consisting of sand-filled EcoFABs, a 17-member SynCom, and the model plant *Brachypodium distachyon*. The EcoFAB is an existing controlled modular growth system designed for reproducible plant-microbiome studies (Gao et al. 2018; Sasse et al. 2019; Zengler et al. 2019). We used quartz sand to provide the root with a physical attachment environment like that in the soil, but minimize the nutritional disturbance to the system (Gao et al. 2018; Sasse et al. 2020). The SynCom we used in this work is a well-characterized microbial community derived from the rhizosphere and adjacent soil of *Panicum virgatum* plants (switchgrass) (Coker et al. 2022). The 17 bacterial strains were selected from different genera, and their 16S rDNA sequences can be readily distinguished. The *B. distachyon* plant was chosen because it is an excellent model to study soil-plant-microbe interactions due to the following advantages. First, it is a monocot model species for cereal and bioenergy crops, as it is phylogenetically closely related to switchgrass, sorghum, wheat, rice, and barley (Draper et al. 2001; International Brachypodium Initiative 2010). *B. distachyon* shares a similar root architecture with economically-important grasses, yet with one-third the size (Hong et al. 2011; Kawasaki et al. 2016; Watt et al. 2009) making it possible to both observe and sample the whole root system in small lab-based experimental systems, such as EcoFABs. Second, it has a small height, rapid life cycle, reduced genome size, and a T-DNA mutant collection is available (Bragg et al. 2012; Draper et al. 2001; Hong et al. 2011; Hsia et al. 2017; International Brachypodium Initiative 2010; Kawasaki et al. 2016; Vogel 2016). Third, the development of *B. distachyon* is well established and standardized using the Biologische Bundesanstalt, Bundessortenamt and CHemische Industrie (BBCH) scale (Hong et al. 2011), which allows easy comparison between experimental setups. Apart from being an excellent grass model for the reasons above, understanding the relationships between *B. distachyon* and microbial communities could reveal novel biomass production enhancement strategies for this plant.

Using the EcoFAB-*B. distachyon*-SynCom set up we evaluated how different SynCom inoculation practices and plant harvesting time points affect the persistence, stability, and consistency of the sand-plant-microbe fabricated ecosystem. Our data showed that for at least 21 days post-inoculation (DPI), the SynCom was highly persistent and stable regardless of inoculation practice, and the plant phenotypes were highly reproducible. The results demonstrate that our set up is a highly conserved system suitable for future plant-microbe interactions studies. This could include, but is not limited to, reproducibility assessment between different labs, resilience studies under biotic and abiotic stress, SynCom strain selection and tailoring, *in situ* ecosystem gene editing, and establishment of more complex fabricated ecosystems that include fungi and/or phage.

## MATERIALS AND METHODS

### EcoFAB preparation

EcoFAB devices (https://eco-fab.org/) were fabricated according to (Gao et al. 2018) with some modifications. Briefly, a siloxane elastomer base-curing agent mixture (1:10 v/v ratio) (polydimethylsiloxane, PDMS; Ellsworth Adhesives, USA) was poured onto a 3D-printed mold and allowed to solidify at 80°C for 4 h. The PDMS layer was separated from the mold, the edges trimmed, the surface cleaned by a plasma cleaner, and permanently bonded to a glass microscope slide by pressing the PDMS and glass slide together. Then EcoFABs were filled with substrates. Hydroponic media consisting of 50%-strength Murashige and Skoog basal medium (0.5X MS, PhytoTech Lab, USA) was used to compare the development of *B. distachyon* grown in EcoFABs with conventional methods. Quartz sand (50-70 mesh particle size, non-acid washed, Sigma-Aldrich, USA) was used for all other experiments. Finally, each device was placed in a GA-7 Magenta box (Fisher scientific, USA) with a customized watering system (Fig. 1) and sterilized by autoclave for 15 min.

**Fig. 1.**
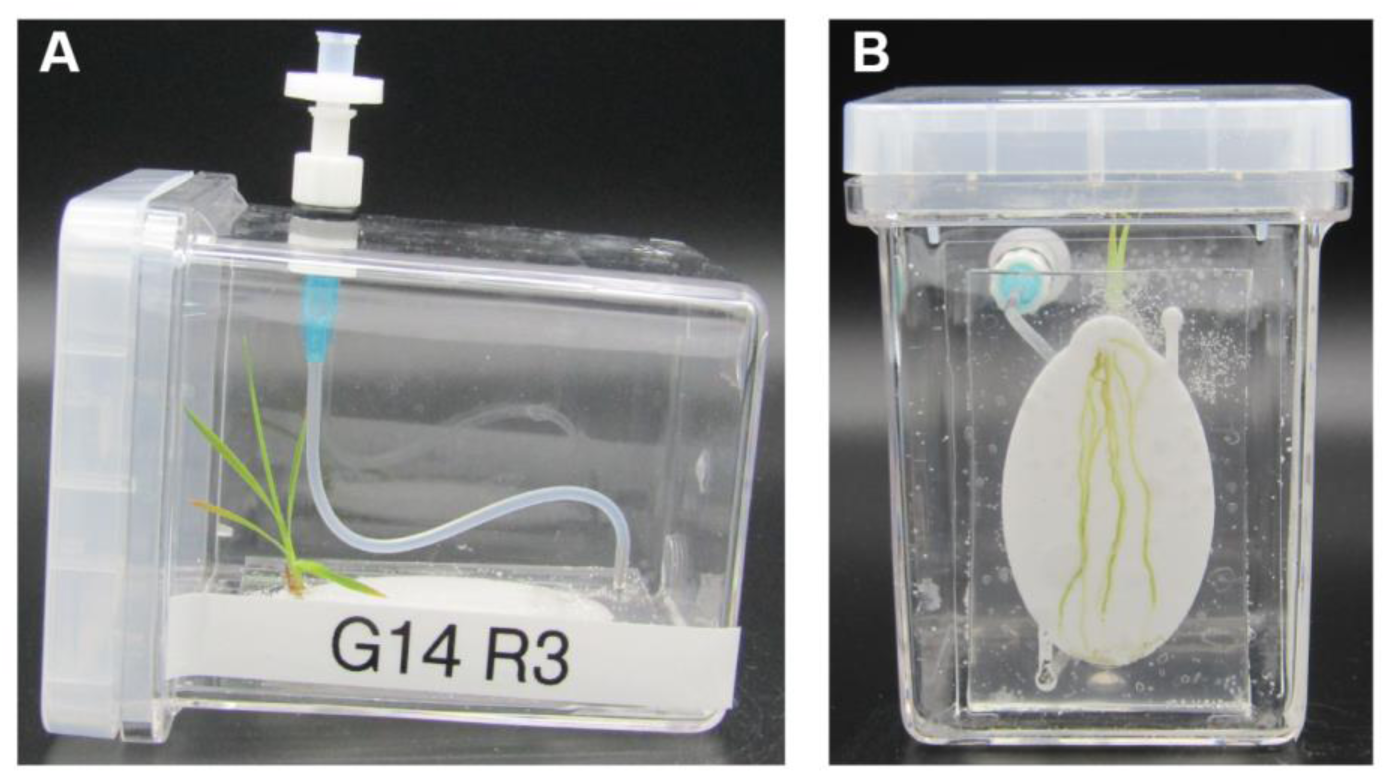
*B. distachyon* grown in a quartz sand-filled EcoFAB inside a Magenta box. **A**, Side view enables shoot observation and **B**, bottom view enables root observation.

### Plant growth conditions

*Brachypodium distachyon* Bd21-3, an inbred diploid line with full genome sequence (Vogel and Hill 2008) (hereafter referred to as Bd21-3), seeds were dehusked and sterilized in 70% v/v ethanol for 2 min, and in 50% v/v NaOCl for 5 min, followed by five wash steps in sterile water. The surface-sterilized seeds were stratified at 4°C in dark for 2 days, and then germinated on sterilized wet filter paper for three days. Bd21-3 seedlings were grown in a growth chamber maintained at 200 µmolm^-2^ s^-1^, 16-hr light/8-hr dark regime, 40% RH, at 24°C. After three days, using sterile forceps, seedlings of comparable size were transferred to EcoFABs with a root growth chamber volume of 3.1 ml. For seeds inoculated at 0 days after germination (0 DAG), sterilized seeds were placed directly into EcoFABs.

After being transferred into an EcoFAB, the root growth chambers were filled with 0.5X MS medium. Each EcoFAB was then placed into a sterile Magenta box (Fisher Scientific, USA), to maintain sterile conditions during the experiment, and maintained in the conditions explained above. EcoFABs were supplemented with sterile water every 2-3 days until harvest through a customized watering system equipped with a 0.22 µm filter (Pall Life Sciences, USA) to avoid opening the system (Fig. 1). When EcoFABs without plants were needed, they were used and incubated in the same conditions as above.

### Bacterial strains and culture conditions

Bacterial strains used in this study (Table 1) were obtained from a *Panicum virgatum* (switchgrass) field in Oklahoma, United States (Coker et al. 2022). Strains were cultured at 30°C with 180 rpm using Reasoner’s 2A (R2A) medium (van der Linde et al. 1999; Reasoner and Geldreich 1985), 10%-strength R2A (0.1X R2A), Luria-Bertani (LB) medium (Thermo Fisher, USA), and Yeast extract-Malt extract (YM) medium (Atlas 2010) as indicated in Table 1. *Bradyrhizobium* OAE829 was prepared by streaking it onto several 0.1X R2A agar plates and growing it at 30°C for 10 days, and then resuspending it in liquid media when needed.

**Table 1.** Bacterial strains of the 17-member SynCom used in this study.

| Strain | Medium |
| --- | --- |
| *Arthrobacter* OAP107 | R2A |
| *Bacillus* OAE603 | R2A |
| *Bosea* OAE506 | R2A |
| *Bradyrhizobium* OAE829 | 0.1X R2A |
| *Brevibacillus* OAP136 | R2A |
| *Burkholderia* OAS925 | R2A |
| *Chitinophaga* OAE865 | R2A |
| *Lysobacter* OAE881 | LB |
| *Marmoricola* OAE513 | R2A |
| *Methylobacterium* OAE515 | LB |
| *Mucilaginibacter* OAE612 | R2A |
| *Mycobacterium* OAE908 | R2A |
| *Niastella* OAS944 | R2A |
| *Paenibacillus* OAE614 | R2A |
| *Rhizobium* OAE497 | YM |
| *Rhodococcus* OAS809 | R2A |
| *Variovorax* OAS795 | R2A |

### Synthetic community preparation and inoculation

Bacterial strains (except *Bradyrhizobium* OAE829) were grown for 48 h in 10 ml of culture media, as explained above. *Bradyrhizobium* OAE829, which does not grow well in liquid medium, was grown on several 0.1X R2A agar plates for 10 days, and the bacterial mass was collected by sterile loops, pooled, and resuspended in 0.5X MS. Cultures were centrifuged at 4,000 x g for 20 minutes and washed twice with 0.5X MS. Then the OD_600_ of each culture was measured and the 17 strains were pooled such that each had a final OD_600_ equal to 1. This SynCom liquid mixture served as the inoculant. The SynCom inoculant (31 µl) was applied directly to each root growth chamber in the EcoFABs. At least 3 biological replicates were used for each condition.

### Sample harvesting

At harvest, samples were collected in sterile conditions unless otherwise specified. The EcoFABs were removed from each Magenta box, the root growth chambers were sliced open with a sterile scalpel, and the plants were taken out. The fresh weight of aerial and root fractions was recorded. Three types of samples were collected: “sand” (bacteria isolated from the sand not attached to the root); “rhizosphere” (bacteria from the sand attached to the roots); and “root” (bacteria from the surface-sterilized root). For the sand sample, the remaining sand in the EcoFAB after the plants were taken out was collected into a 50 ml tube containing 25 ml phosphate buffered saline (PBS; pH 7.2; Thermo Fisher Scientific, USA). For the rhizosphere sample, the sand covering the root was obtained by placing the root into a 50 ml tube containing 25 ml PBS, and vortexed for 10 seconds, twice. To obtain the root sample, the root was surface sterilized by washing three times with Milli-Q water (Millipore, USA), then with 3% v/v H_2_O_2_ for 30 seconds, followed by a wash with 100% v/v EtOH for 30 seconds, 6.15% v/v NaClO with 0.01% Tween-20 for 3 minutes, 3% v/v H_2_O_2_ for 30 seconds, and five final washes with sterile Milli-Q water for 30 seconds. In the case of EcoFABs without plants, only the “sand” sample was harvested. Samples were stored at -80°C until processed for bacterial DNA extraction.

### DNA extraction and 16S rDNA amplicon sequencing

The DNeasy PowerSoil HTP 96 Kit (Qiagen, USA) was used to extract bacterial DNA according to the manufacturer’s manual. Sand and rhizosphere samples were prepared by adding 250 mg of the collected substrate to a PowerBead plate and homogenized using a TissueLyser II (Qiagen, USA) at a frequency of 20 Hz for 10 minutes, twice. Root samples were homogenized by a ¼’’ precision ball using a TissueLyser II (Qiagen, USA) at 30 Hz for 3 minutes, twice. Homogenized root samples were transferred to a PowerBead plate and DNA was extracted with the other samples.

DNA concentrations were quantified by a Quant-iT dsDNA High-Sensitivity kit (Thermo Fisher Scientific, USA) and normalized to 0.3 ng/µl. DNA template was added to a PCR reaction to amplify the V4/V5 16S gene region using the 515F/926R primers, based on the Earth Microbiome Project primers (Parada et al. 2016; Quince et al. 2011) but with in-line dual Illumina indexes (Price et al. 2018; Sharpless et al. 2022). The amplicons were sequenced on an lllumina MiSeq with 600 bp v3 reagents (Illumina, USA).

### Data analysis and visualization

The 16S rRNA gene amplicons were analyzed as follows. Reads were processed with custom Perl scripts implementing Pear for read merging (Zhang et al. 2014). USearch (Edgar 2010) was used to map reads to a database of SynCom V4/V5 16S region sequences using the ‘annot’ command (Edgar 2016) and identifying and removing chimeras. The RDP database (Cole et al. 2014) was used to assign taxonomy to unidentified amplicons. Data visualization was performed within Jupyter notebooks (6.3.0) using either the Seaborn (0.11.1) and matplotlib (3.3.4) packages in Python (3.8.8) or within RStudio (v2023.03.0; RStudio Team) using the ggplot2 (3.3.6) package in R (v4.2.2).

Statistical analyses were performed within RStudio (v2023.03.0; RStudio Team) using R (v4.2.2). Statistical analyses on plant growth were performed using one-way ANOVA followed by Tukey’s HSD test (alpha = 0.05). Shannon diversity index was calculated using the vegan (2.6.4) package and statistical analysis was performed using one-way ANOVA followed by Tukey’s HSD test (alpha = 0.05). Beta diversity was calculated by Bray-Curtis distance using vegan (2.6.4), and permutational multivariate analysis of variance (PERMANOVA) was calculated using adonis2 in the vegan (2.6.4) package. Pairwise comparisons were performed using DEseq2 (1.38.3). Reproducibility was calculated using the within group Euclidean distances among replicates described in (Song et al. 2021).

## RESULTS

### The developmental stages of *B. distachyon* grown in an EcoFAB aligns with those of conventional methods

To compare the development of *B. distachyon* grown in EcoFAB with the growth stages observed using conventional plant growth methods (Hong et al. 2011), we used hydroponic media (0.5X MS) instead of quartz sand in order to better visualize root development. After 3 days in the light, the seeds had germinated and had a visible radicle and coleoptile (Fig. 2), corresponding to day 2.9 in (Hong et al. 2011) (Supplemental Table S1). We then transferred the 3-day-old seedlings into hydroponic EcoFABs to observe their growth over time. Eight days after germination (DAG) the first leaf of the seedling had unfolded and two coleoptile node roots had appeared (Fig. 2), which was observed at day 7.6 in (Hong et al. 2011) (Supplemental Table S1). On day 14, the shoot had 3 unfolded true leaves and the first tiller became detectable, and the root started to reach the far end of the EcoFAB root growth chamber (Fig. 2). The “3 unfolded true leaves” stage was observed at day 15.0 in (Hong et al. 2011). Both shoot and root developed quickly after day 14, and the root system occupied most of the root chamber at day 21 (Fig. 2). On day 28 DAG, the plant began to enter the reproductive phase with an extending flag leaf sheath, and the root system continued developing and occupied the root growth chamber (Fig. 2), corresponding to day 24.9 in (Hong et al. 2011) (Supplemental Table S1).

**Fig. 2.**
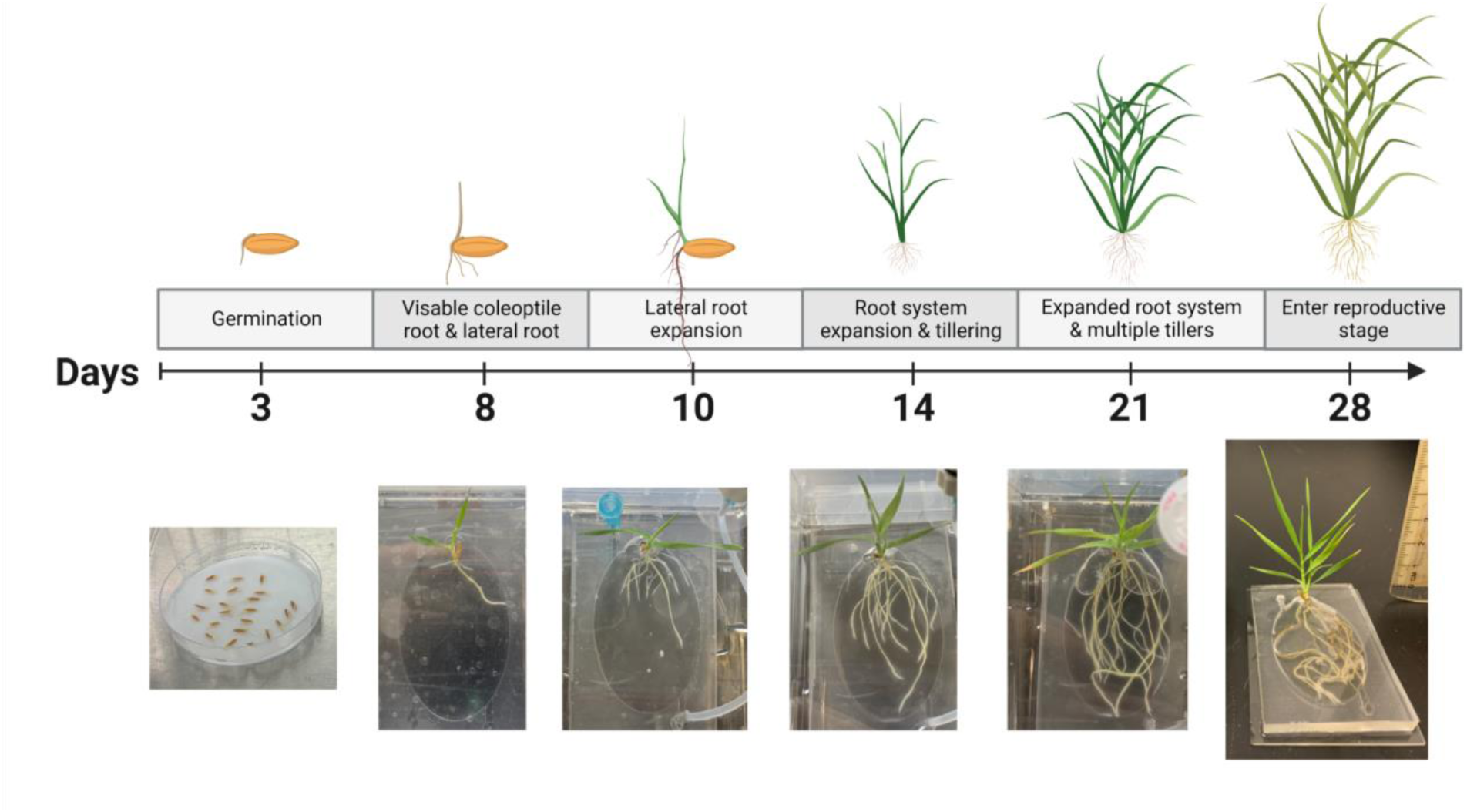
Overview of *B. distachyon* growth in hydroponic EcoFABs. *B. distachyon* seeds were germinated on wet filter paper for three days, and then moved to EcoFABs containing 50%-strength Murashige and Skoog (0.5X MS) plant media. Coleoptile root and lateral root formation happened 8 days after germination (DAG). Formation of the first tiller and expansion of the root system to the bottom side of the EcoFAB root growth chamber happened at 14 DAG. At around 21 DAG, the root system expanded and occupied the whole root growth chamber. The plant entered the reproductive phase around 28 DAG, when most of the EcoFAB chamber was occupied by roots.

### In the absence of a plant, *Variovorax* dominates the microbial community in the EcoFAB

To understand how the SynCom community assembled temporally without a plant, we inoculated SynCom to sand-filled EcoFABs and harvested the samples at 0, 6, 11, 14, and 21 DPI. At each harvest point, the sand samples were collected and relative abundance of the SynCom members was determined by 16S rDNA sequencing. As a control, the liquid inoculant that was used for inoculating the EcoFAB was also collected.

The results show that SynCom composition of the liquid inoculant was similar to that of the one recovered from sand at 0 DPI (Fig. 3A). *Variovorax* OAS795 dominated the population as early as 6 DPI, and remained a dominant species (> 55% of the population) until 21 DPI (Fig. 3A). In contrast, the abundance of *Lysobacter* OAE881, *Niastella* OAS944, and *Bacillus* OAE603 significantly decreased to lower than 0.01% from 6 DPI and onward, respectively. *Brevibacillus* OAP136 and *Bradyrhizobium* OAE829 were not detected at 6 DPI and thereafter. *Marmoricola* OAE513 and *Mycobacterium* OAE908 could not be detected in the liquid inoculant, despite being included in the SynCom at the same ratio as other bacterial strains based on OD_600_. To explore the impact of co-culture timing on bacterial composition, we performed a Principal Coordinate Analysis (PCoA) of Bray-Curtis dissimilarity on all samples. The samples clustered primarily by DPI (PERMANOVA, *F* = 8.3, *P* = 0.0007) and by sample type (PERMANOVA, *F* = 4.8, *P* = 0.0232), which together explained 77.6% of the variation (Fig. 3B). The dissimilarity within groups was smaller when the co-culture time increased and was most similar between 14 DPI and 21 DPI. These results imply that the SynCom in quartz sand in the absence of plants reached a stable community from 14 DPI onward.

**Fig. 3.**
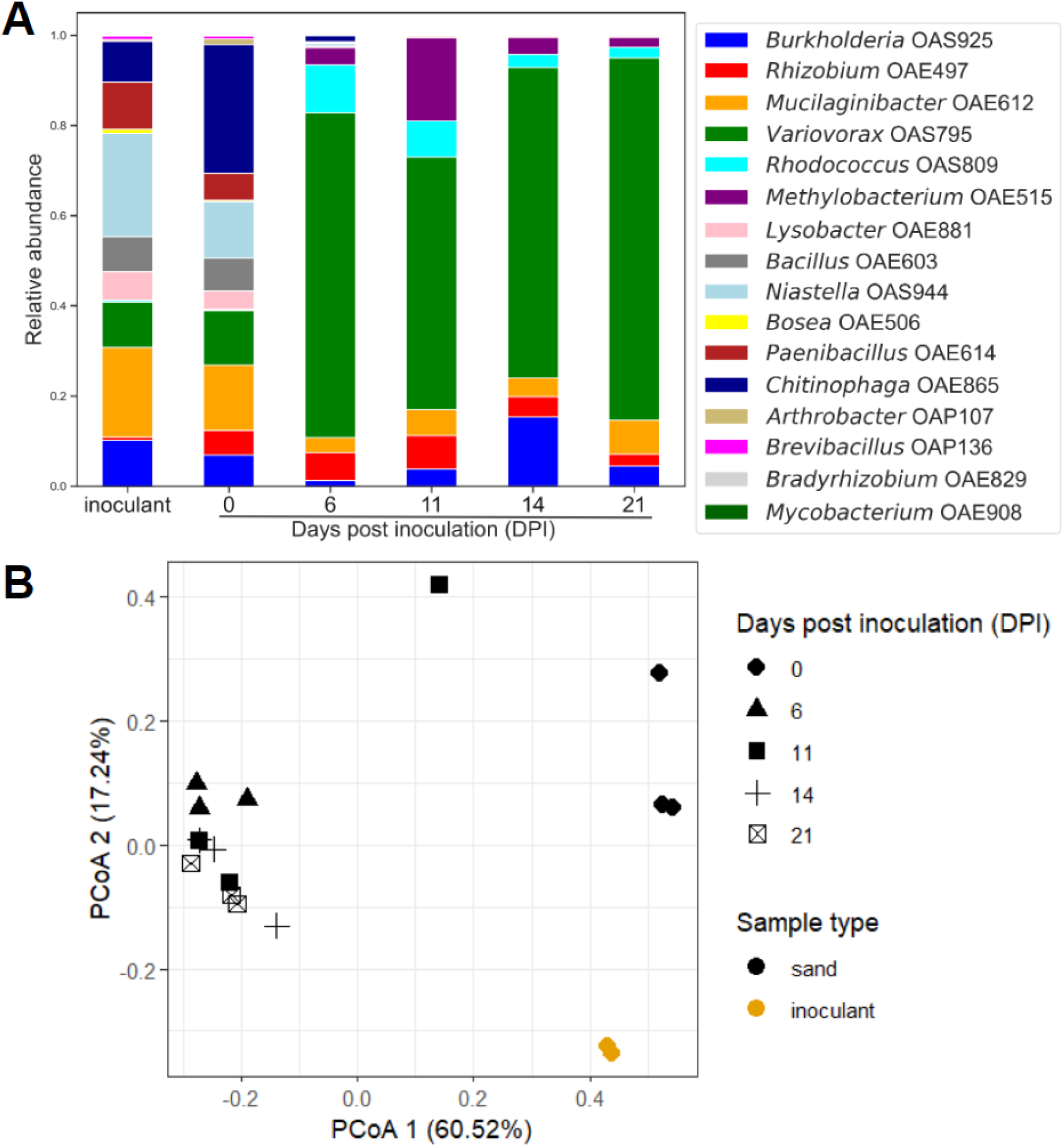
Abundance of SynCom members in quartz sand-filled EcoFABs in the absence of a plant. The microbial community recovered from SynCom liquid inoculant (represented as “inoculant”) and from sand-filled EcoFABs at different inoculation timing. **A**, Mean relative abundance of each bacterial species over time. The SynCom composition of the liquid inoculant was similar to the composition of the microbiome recovered from sand at 0 days post-inoculation (DPI). At 6 DPI in the sand-filled EcoFABs, *Variovorax* OAS795 consisted of more than 50% of the total population while *Lysobacter* OAE881, *Niastella* OAS944, and *Bacillus* OAE603 significantly decreased to lower than 0.01%, respectively. This trend persisted until 21 DPI. **B**, Between-sample diversity analysis using Principal Coordinate Analysis (PCoA). SynCom liquid inoculant used to inoculate the EcoFAB is shown in yellow while the samples recovered from sand-filled EcoFABs are shown in black. Days post-inoculation (DPI) is shown with different shapes. Samples clustered primarily by DPI (PERMANOVA, *F* = 8.3, *P* = 0.0007) and sample type (PERMANOVA, *F* = 4.8, *P* = 0.0232), which together explained 77.6% of the variation. Two and three biological replicates were used respectively for the liquid inoculant and sand-EcoFABs.

### The rhizosphere microbiome is significantly affected by inoculation practices

To assess how inoculation practices shaped microbial communities in the presence of the plant, we inoculated *B. distachyon* with the SynCom using one of three methods: seed inoculation (inoculation at 0 DAG; hereafter indicated as ‘’0 DAG’’), transplant inoculation (inoculation at 3 DAG in parallel with transferring the plant into an EcoFAB), and seedling inoculation (inoculation at 8 DAG when coleoptile node roots and lateral roots started to develop) (Supplemental Fig. S1). Microbiome community samples from sand, rhizosphere, and root were collected when *B. distachyon* reached 14 DAG (Fig. 3). This enabled us to elucidate the effects of inoculation practices regardless of the plant developmental stage.

The 14 DAG *B. distachyon* plants had at least 3 unfolded true leaves and the root had reached the far end of the EcoFAB regardless of the SynCom inoculation practice (Supplemental Fig. S2), with a similar morphology to that observed in hydroponic EcoFABs (Fig. 3). There were no significant differences in fresh weight between different inoculation practices (ANOVA, *P* = 0.406 for shoot and *P* = 0.134 for root, Supplemental Fig. S3). We determined the within-sample diversity (alpha diversity) of the SynCom using the Shannon diversity index (Fig. 4A). The results showed that all root samples had significantly lower alpha diversity than their rhizosphere and sand counterparts (ANOVA, *P* = 1 x 10^-10^), and that different inoculation practices had similar alpha diversity (ANOVA, *P* > 0.05) in each sample type (Fig. 4A).

**Fig. 4.**
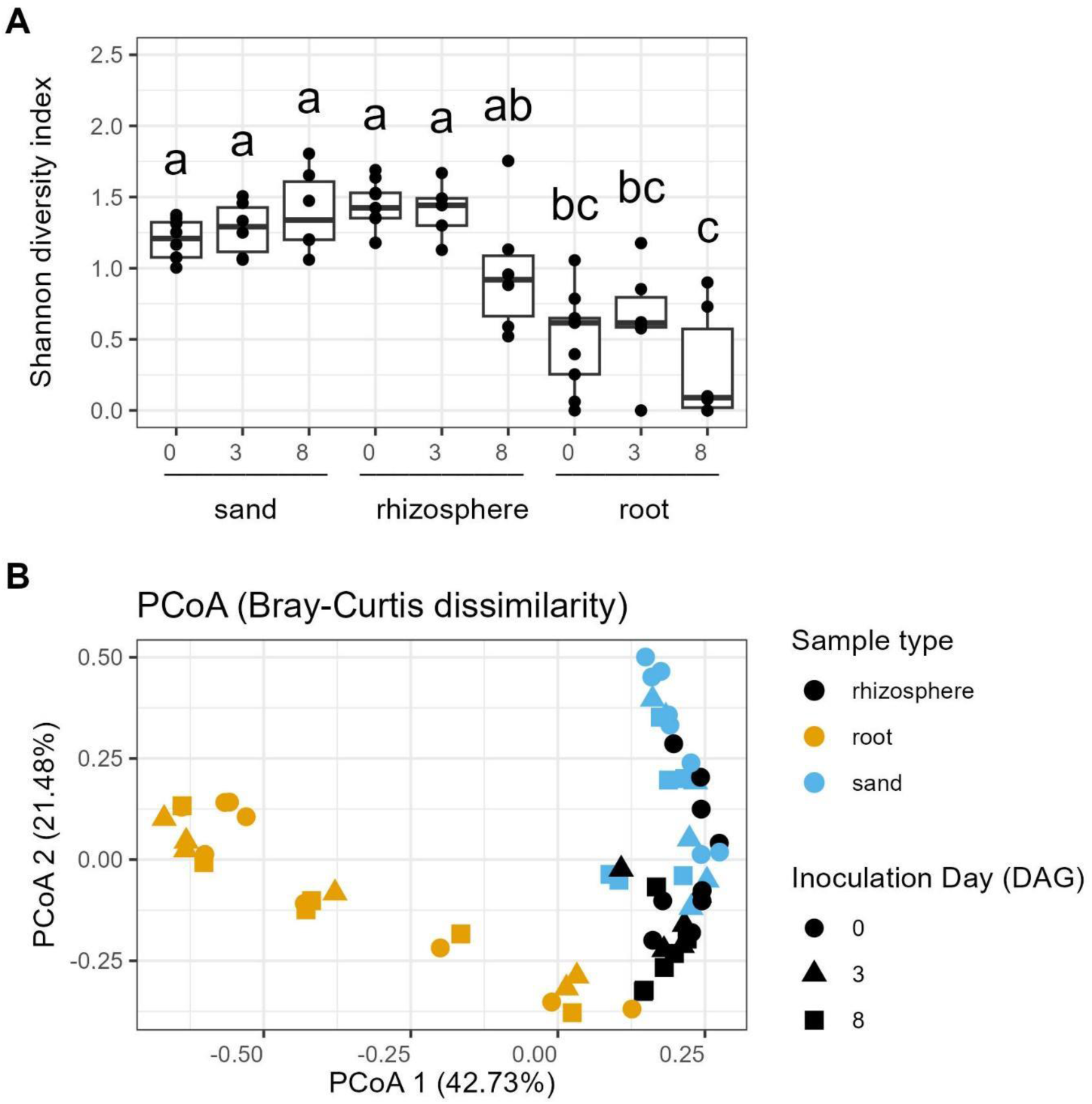
Effect of both sample type and inoculation practices in the microbial community of *B. distachyon* plants 14 days after germination (DAG). The microbial community of the tillering *B. distachyon* plants was significantly affected by both sample type and by inoculation practices. The SynCom inoculation was performed at 0, 3, and 8 DAG and the microbiome samples from sand, rhizosphere, and root were collected when the plant entered the tillering stage (14 DAG). **A**, Boxplot of Shannon’s diversity index of sand, rhizosphere, and root (*P* = 1.0 x 10^-10^). The root samples had a significantly lower Shannon’s diversity index than the rhizosphere and sand ones. The Shannon’s diversity index was not significantly different with respect to sample type (sand, rhizosphere, and root). Statistical analysis consisted of ANOVA followed by Tukey HSD (alpha = 0.05). Values that do not share a common letter label are significantly different from each other. **B**, Principal Coordinate Analysis (PCoA) of all amplicons across different SynCom inoculation practices and sample types. Samples were colored by sample type and shaped according to microbiome inoculation practice. A PERMANOVA analysis showed that the diversity of the samples was best explained by sample type (*F* = 21.9, *P* < 0.0001) and moderately by inoculation practice (*F* = 1.9218, *P* = 0.058). At least 6 biological replicates were used in each group.

To further explore the between-sample diversity (beta diversity), we performed a PCoA test of Bray-Curtis dissimilarity. The PCoA1 (42.73%) was best explained by the variation within the root samples, and PCoA2 (21.41%) clustered among sand and rhizosphere samples (Fig. 4B). The top two (PCoA1 plus PCoA2) axes explained 61.41% of the variance. A PERMANOVA analysis showed that beta diversity was best explained by sample type (*F* = 21.9, *P* < 0.0001) and moderately by inoculation practice (*F* = 1.9218, *P* = 0.058). To further evaluate whether different inoculation practices affected microbiome assembly within each sample type, we performed a PERMANOVA test among different inoculation practices. The results indicated that the microbial community was significantly affected by inoculation practice in the rhizosphere samples (*P* = 0.0059), moderately affected in sand samples (*P* = 0.08309), but not affected in root samples (*P* = 0.7154) (Supplemental Table S2).

To gain a better understanding of how inoculation practices affected the rhizosphere microbial community, we performed pairwise comparisons between inoculation practices for each SynCom member (Fig. 5). The rhizosphere *Muciliaginibacter* OAE612 in the 3 DAG group was significantly lower than that of the 0 DAG group (DESeq2, Benjamini– Hochberg corrected *P* = 0.0059, Fig. 5A). When comparing 8 DAG rhizosphere microbial communities with that of 0 DAG group, *Burkholderia* OAS925 and *Lysobacter* OAE881 were enriched (DESeq2, Benjamini–Hochberg corrected *P* = 0.0009 and 0.0001, respectively, Fig. 5B) while *Variovorax* OAS795 decreased (DESeq2, Benjamini– Hochberg corrected *P* = 0.0008, Fig. 5B). When comparing 8 DAG and 3 DAG groups, *Burkholderia* OAS925 and *Lysobacter* OAE881 were enriched (DESeq2, Benjamini– Hochberg corrected *P* = 0.0240 and 0.0092, respectively, Fig. 5B) while *Methylobacterium* OAE515 and *Rhodococcus* OAS809 decreased (DESeq2, Benjamini– Hochberg corrected *P* = 0.0063 and 0.0072, respectively, Fig. 5B). In summary, the data demonstrated that inoculation practices significantly affected the rhizosphere microbial communities of 14 DAG plants.

**Fig. 5.**
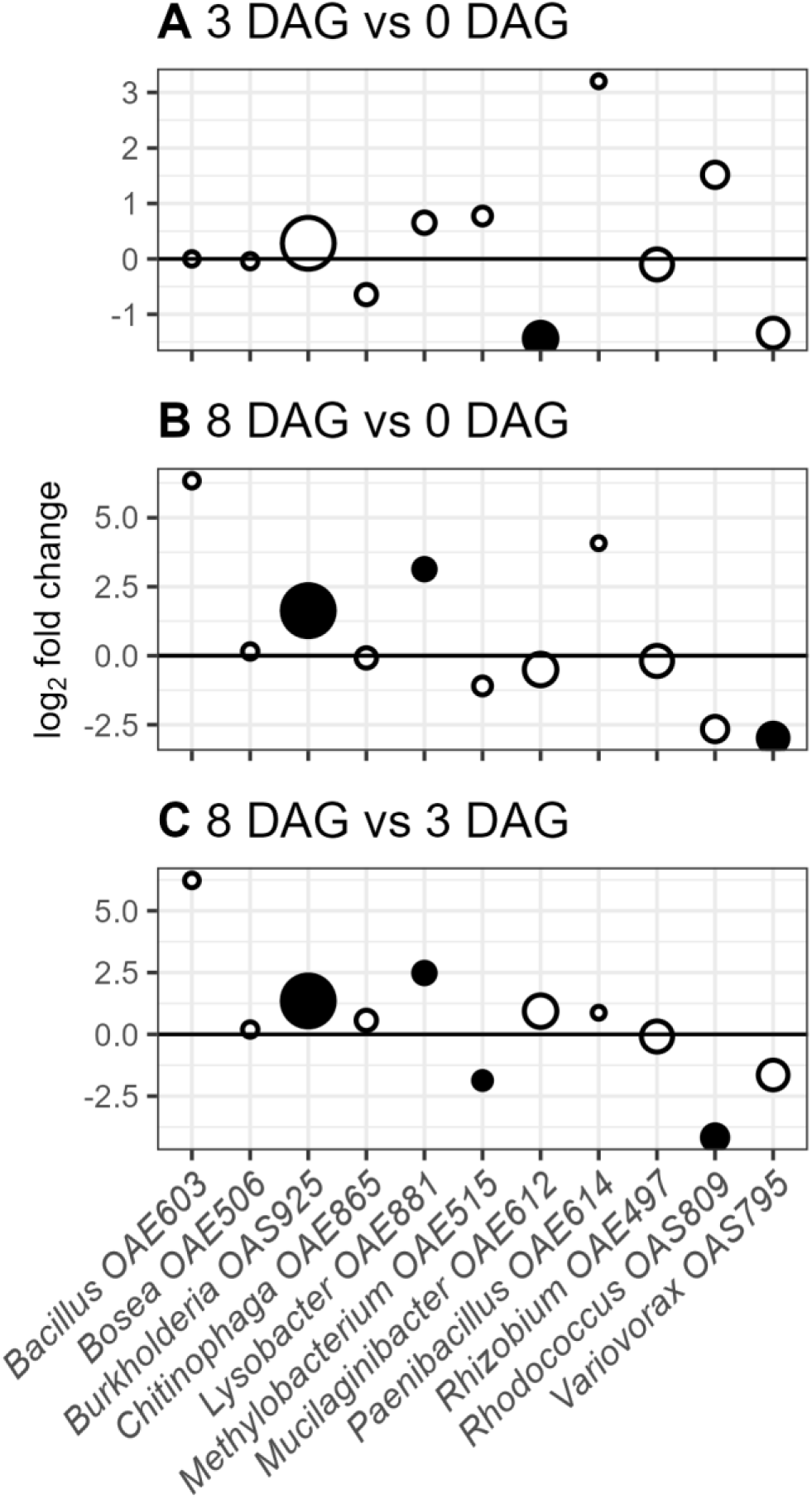
Pairwise comparison of inoculation methods in *B. distachyon* rhizosphere microbial communities 14 days after germination (DAG). DESeq2 analysis revealed the differentially abundant bacteria. **A**, Abundance of *Mucilaginibacter* OAE612 was decreased in 3 DAG compared to 0 DAG. **B**, *Burkholderia* OAS925 and *Lysobacter* OAE881 were enriched in 8 DAG compared to 0 DAG, while *Variovorax* OAS795 was depleted. **C**, When comparing 8 DAG and 3 DAG, *Burkholderia* OAS925 and *Lysobacter* OAE881 were enriched while *Methylobacterium* OAE515 and *Rhodococcus* OAS809 were depleted. The y axis represents the log_2_ fold change and the positive values indicate higher abundance in the former groups. Differentially abundant bacteria (*Padj* < 0.05) are shown in filled circles and others are shown in open circles. Circle size represents the base mean. Statistical analysis was performed using DESeq2 with default settings. At least 6 biological replicates were used in each group.

### The transplant inoculation is most reproducible when the microbial community reaches a steady state

To further understand whether different inoculation practices would result in a similar, steady-state microbiome in the different sample types, we inoculated the SynCom using seed inoculation (0 DAG), transplant inoculation (3 DAG), and seedling inoculation methods (8 DAG). Samples were harvested at 21 DPI (Supplemental Fig. S4). This prolonged co-culture setting enabled us to test the persistence and stability of the SynCom in association with the plant. Samples of sand, rhizosphere, and root were collected and resolved by 16S rRNA gene sequencing. At 21 DPI (corresponding to 21 DAG, 24 DAG and 29 DAG plants), *B. distachyon* leaves had brown leaf tips in every group, and the shoot of the 21 DAG group plants showed a smaller size 29 DAG group plants (Supplemental Fig. S6). The root system had occupied most of the root growth chamber (Supplemental Fig. S6), similar to what was observed in hydroponic EcoFABs (Fig. 2).

For each inoculation method, the Shannon diversity index of the root samples was significantly lower than that of the rhizosphere and sand samples (Fig. 6A). Among the root samples, the Shannon diversity index of the seed inoculation group was significantly higher than that of the other inoculation methods (Fig. 6A); this phenomenon was not observed in the rhizosphere or sand samples. To explore whether the different inoculation practices result in microbial composition differences, we analyzed their beta diversity by PCoA test using Bray-Curtis dissimilarity (Fig. 6B). The top two axes (PCoA1 and PCoA2) explained 68.73% of the variance, and the samples were separated by their sample type (PERMANOVA, *F* = 34.25, *P* < 0.0001; Supplemental Table S3) but not by the inoculation practice (PERMANOVA, *F* = 1.5436, *P* = 0.1469). These results indicated that when reaching a steady state, the microbial communities were very similar within-sample types, regardless of their inoculation practice.

**Fig. 6.**
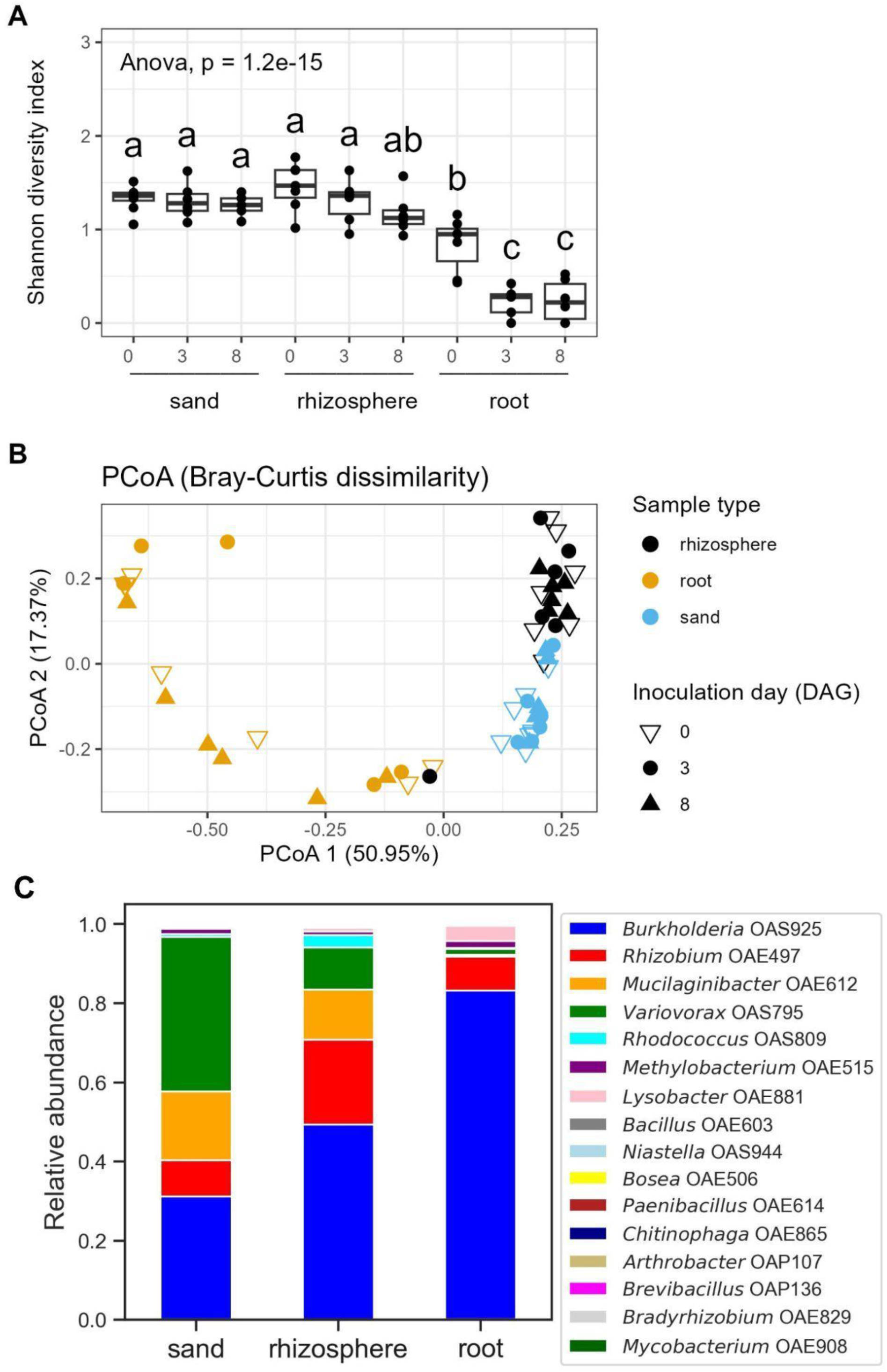
Effect of both sample type and inoculation practices on the microbial community of *B. distachyon* plants 21 days post-inoculation (DPI). The microbial community at 21 DPI was best explained by their proximity to *B. distachyon* roots rather than inoculation day. The SynCom inoculation was performed at 0, 3, and 8 days after germination (DAG) and the microbiome samples from sand, rhizosphere, and root were collected at 21 DPI. **A**, Boxplot of Shannon’s diversity index of sand, rhizosphere, and root (*P* = 1.2 x 10^-15^). **B**, Principal Coordinate Analysis (PCoA) of all amplicons across different SynCom inoculation time and sample types. **C**, Relative abundance of each strain per sample type at 21 DPI. All data from the same sample type were merged and the relative abundance was calculated. At least 6 biological replicates were used in each group in A and B. Biological replicates of each group in C were equal or larger than 17.

To assess how sample type shapes the microbiome, we combined all the data from each sample type and plotted the relative abundance of the bacterial species. The results showed that the relative abundance of *Burkholderia* OAS925 was positively correlated with the proximity to the plant, while the relative abundance of *Variovorax* OAS795 and *Mucilaginibacter* OAE612 were negatively correlated (Fig. 6C). While all bacteria detected in the liquid inoculant were detected in the rhizosphere at 21 DPI, *Bradyrhizobium* OAE829 and *Paenibacillus* OAE614 were not detected in the sand. Regarding the root samples, *Bradyrhizobium* OAE829, *Paenibacillus* OAE614, *Chitinophaga* OAE865, *Brevibacillus* OAP136, and *Niastella* OAS944 were not detected (Fig. 6C). These results indicated that different bacteria established distinct niches in the steady state microbial community.

To better understand how the plant roots shape their microbial community, we performed pairwise comparisons on bacterial abundance between rhizosphere and sand, as well as between root and rhizosphere (Fig. 7). The results showed that *Burkholderia* OAS925 was significantly enriched in the rhizosphere compared to sand, while *Variovorax* OAS795 and *Mucilaginibacter* OAE612 were significantly diminished (DESeq2, Benjamini–Hochberg corrected, *P* = 0.0329, 5.052 x 10^-14^ and 0.0152, respectively, Fig. 7A). In addition to *Burkholderia* OAS925, *Rhizobium* OAE497, *Lysobacter* OAE881, and *Rhodococcus* OAS809 were significantly enriched in the rhizosphere as compared to the sand (*P* = 2.3561 x 10^-11^, 0.0152, and 0.0021, respectively, DESeq2, Benjamini– Hochberg corrected, Fig. 7A). These results agree with our observations on the merged relative abundance (Fig. 6C). When comparing between the root and the rhizosphere, *Mucilaginibacter* OAE612, *Variovorax* OAS795, *Rhodococcus* OAS809, and *Rhizobium* OAE497 were significantly diminished in the root compared to in the rhizosphere (DESeq2, Benjamini–Hochberg corrected, *P* = 1.180 x 10^-21^, 1.7171 x 10^-8^, 3.9037 x 10^-8^, 7.9962 x 10^-4^, respectively, Fig. 7B). The results suggested the root is a more selective niche than the rhizosphere.

**Fig. 7.**
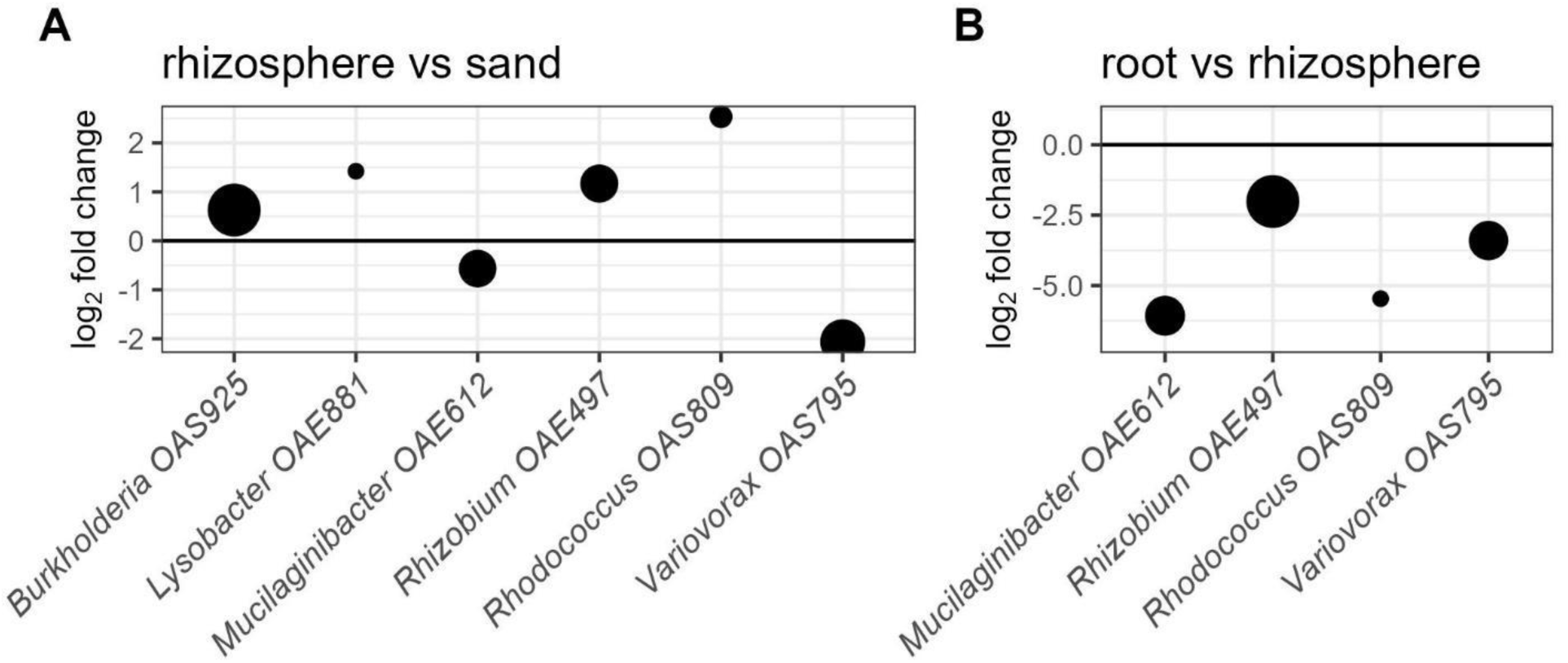
Pairwise comparison of the microbial communities collected in *B. distachyon* EcoFABs 21 days post-inoculation (DPI). **A**, The DESeq2 analysis revealed the differentially abundant bacteria between the rhizosphere and sand. The x axis represents the log_2_ fold change and the positive values indicate higher abundance in the rhizosphere. **B**, The DESeq2 analysis revealed the differentially abundant bacteria between root and rhizosphere. The x axis represents the log_2_ fold change and the positive values indicate higher abundance in the root. Differentially abundant bacteria (*Padj* < 0.05) are shown. Circle size represents the base mean. Statistical analysis was performed using DESeq2 by a Wald test and was corrected using the Benjamini–Hochberg method for multiple testing. Biological replicates of each group in C were equal or larger than 17.

To identify the best inoculation practice, we evaluated how inoculation methods impacted the plant fresh weight and microbial community reproducibility. We collected the root and shoot fresh weight at 21 DPI. Three uninoculated plants were added to each group to serve as a control. The SynCom inoculation had no significant effect on plant biomass except for the root weight of the 8 DAG group, which was negatively affected by SynCom inoculation (t-test, *P* = 0.0140; Supplemental Fig. 7). We then asked which inoculation method has the highest reproducible microbiome community within a group. We calculated the microbial community reproducibility using Euclidean distance as described previously (Song et al. 2021), where the Euclidean distance was negatively correlated with the reproducibility. These results show that there were no significant differences between the sand and root samples (t-test, *P* = 0.074 and 0.160, respectively; Fig. 8A and 8C), indicating the reproducibility of the sand and root samples were not significantly different between inoculation practices. However, in rhizosphere samples, the 3 DAG group had the lowest median and was significantly lower than the 0 DAG group (t-test, *P* = 0.0046; Fig. 8B). Therefore, the 3 DAG group had the highest reproducibility.

**Fig. 8.**
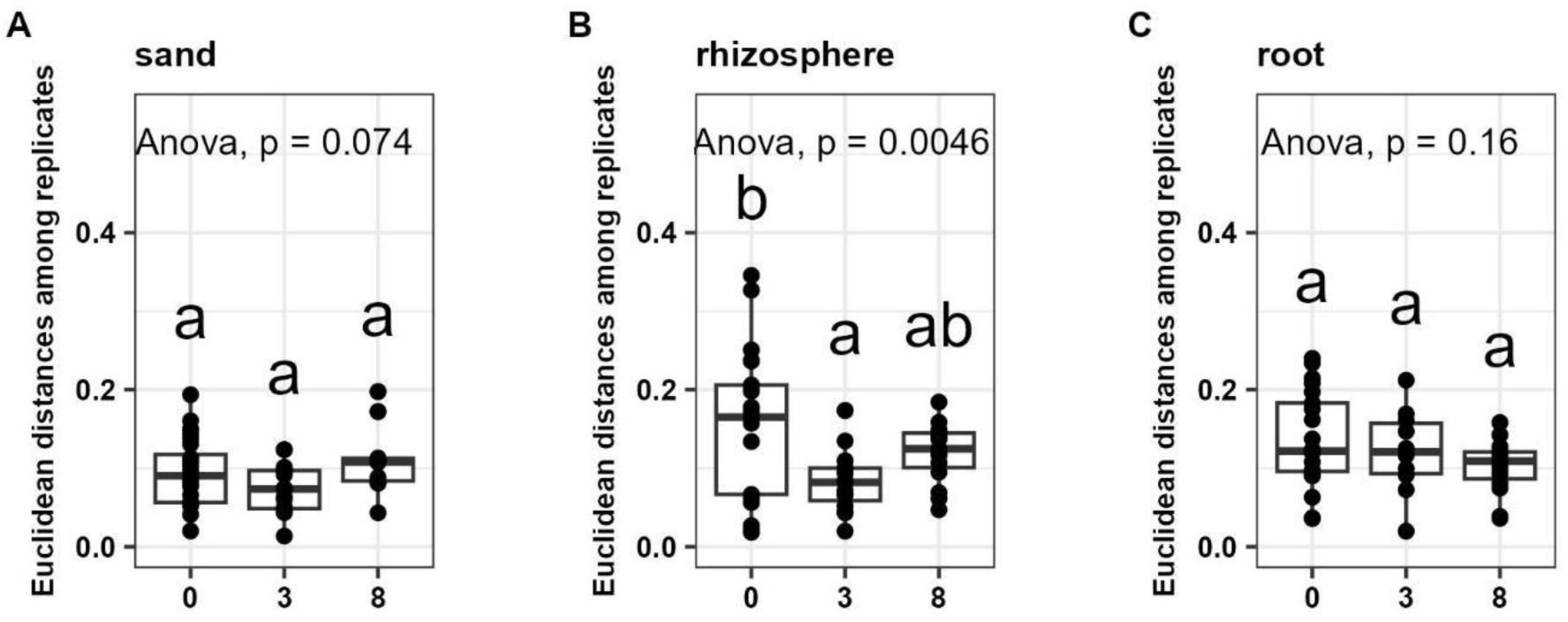
Correlation of the Euclidean distance among replicates and the different inoculation practices (0, 3, and 8 days after germination; DAG). The reproducibility of the microbial community between different inoculation practices is negatively correlated with the Euclidean distance among replicates. **A**, sand, **B**, rhizosphere, and **C**, root samples harvested 21 days post-inoculation (DPI). The reproducibility of the sand and root samples were not significantly different between inoculation practices. In the rhizosphere, the 3 DAG group had the lowest Euclidean distance among replicates and was significantly lower than the 0 DAG group, indicating that the 3 DAG group had the highest reproducibility. Biological replicates of each group in C were equal or larger than 17.

## DISCUSSION

Synthetic microbial communities (SynComs) are a valuable tool for exploring the complexity of interactions between microbes, plants, and the environment. In the last decade, numerous studies have given more insight into rhizosphere dynamics and structure using SynComs. However, more systematic and standardized methodologies are needed to harness the full potential of SynCom tools. A systematic review of the use of SynComs to understand the relationship between plants, microbes and the environment was recently performed (Marín et al. 2021). The authors found that most SynCom studies published so far use *Arabidopsis thaliana* as a plant model, with far fewer using economically-important food or biofuel crop models, such as *Brachypodium distachyon*. Additionally, little standardization and comparison between protocols was observed in the methodologies (Marín et al. 2021). Most SynCom studies focus on microbial abundance composition and *in planta* effect. Studies assessing the impact of inoculation method, plant age, or harvesting point in the microbial community are scarce.

Our current research provides a baseline and foundational information for using the EcoFAB - *B. distachyon* - SynCom as a model fabricated ecosystem for plant-microbe interactions studies, with a particular focus on how inoculation practices impact microbiota assembly and community stability in a small-scale fabricated ecosystem. The ideal system has persistent plant-microbe associations, stable microbial community structure, and produces reproducible plant phenotypes. Using EcoFABs, we tested how harvesting time points (14 DAG, or 21 DPI) and different inoculation methods (seed inoculation-0 DAG, transplant inoculation-3 DAG, and seedling inoculation-8 DAG) affect the microbial community assembled.

Our data showed that all bacteria from the initial inoculant were detected in the rhizosphere samples at 21 DPI, demonstrating that the SynCom had a persistent association with *B. distachyon* for 21 DPI, regardless of inoculation methods. Moreover, the microbial community of each sample type (i.e. sand, rhizosphere, root) was indistinguishable between inoculation practices. Among the samples collected at 21 DPI, the transplant inoculation (3 DAG group) showed the most reproducible microbial community and a neutral plant growth effect. Therefore, we concluded that transplant inoculation is the best inoculation practice to study the steady state microbiome in our system.

### The sand-filled EcoFAB-*B. distachyon*-SynCom is a suitable bench-top fabricated ecosystem model

The results obtained in this study show that the sand-filled EcoFAB-*B. distachyon*-SynCom system aligns with conventional plant growing methods and soil microbial community assembly data. On the plant side, we showed that the developmental stage of *B. distachyon* grown in an EcoFAB (Fig. 2) corresponds well to the conventional growth method documented by (Hong et al. 2011) (Supplemental Fig. S1). When comparing the *B. distachyon* plants in sand-filled EcoFABs to the ones in the hydroponic set up, they both had 3 unfolded leaves at 14 DAG and a flag leaf sheath extending at 28 DAG (Fig. 2, Supplemental Fig. S4, Supplemental Fig. S5). This demonstrated that the developmental stages of *B. distachyon* grown in a sand-filled EcoFAB aligned well with both hydroponic EcoFABs and conventional growth methods.

On the microbial side, this study demonstrated that the SynCom formed a persistent plant-microbe assembly for at least 21 DPI. Our data showed that *Burkholderia* OAS925 (order *Burkholderiales*) and *Lysobacter* OAE881 (order *Xanthomonadales*) were enriched in rhizosphere samples at 21 DPI (Fig. 7). Our results agree with the observations that *Burkholderiales* and *Xanthomonadales* are enriched in *Brachypodium* rhizosphere microbial communities collected from agricultural soil (Kawasaki et al. 2016). One reason why these two orders are usually abundant in the plant environment is because they are fast growing and metabolically adaptable bacteria. Our data is also in line with a recent publication using the same SynCom in hydroponic EcoFABs, where it was shown that *Burkholderia* OAS925 dominated the rhizosphere microbial community 7 DPI (Coker et al. 2022). Coker et al. also showed that when the SynCom were mixed at ratios 1:1 based on their OD_600_, *Marmoricola* OAE513 and *Mycobacterium* OAE908 had very low relative abundance and both wasn’t detectable 3 days after SynCom mixing. This may explain why both bacteria were not detected in our initial inoculant.

### The presence of *B. distachyon* plants influences the composition of the microbiome

The data obtained in this work shows that, in the absence of a plant, *Variovorax* OAS795 (order *Burkholderiales*) dominated the population (> 55% of the population) (Fig. 3A). In the absence of *B. distachyon*, the sand is an environment poor in carbon sources, so the ability of the *Variovorax* strain to outcompete the rest of syncom members must rely on its metabolic adaptability. Members of the genus *Variovorax* have indeed been described as metabolically versatile: it has been shown that *Variovorax* possess extraordinary degradation abilities and can utilize natural compounds produced by other bacteria, such as acyl homoserine lactones (AHLs), as carbon sources (Leadbetter and Greenberg 2000). Additionally, genome analysis of several strains revealed metabolic features for autotrophic lifestyle in some strains of this species (Han et al. 2011).

In the presence of a plant, *Burkholderia* OAS925 dominated the population in the rhizosphere and the root (Fig. 6C). *Burkholderia* species are reported among the dominant bacteria in the rhizosphere of several plants, including *B. distachyon*, wheat (*Triticum aestivum*), maize (*Zea mays*), and sorghum (*Sorghum bicolor*) (Donn et al. 2015; Hallmann et al. 1999; Kawasaki et al. 2016; Li et al. 2014; Parke and Gurian-Sherman 2001; Schlemper et al. 2017). The concentration of *Burkholderia* species in plant roots has been estimated to be 100 times higher than that in bulk soil (Pallud et al. 2001). Their versatile metabolism and adaptability allow them to degrade different compounds usually found in root exudates, such as organic acids, sugars, and polyalcohols (Compant et al. 2008; Morya et al. 2020).

To better understand how plant roots shape the microbial community, we performed pairwise comparisons between rhizosphere and sand, as well as between root and rhizosphere. The results showed that *Burkholderia* OAS925 was significantly more enriched in the rhizosphere than in the sand, while *Variovorax* OAS795 and *Mucilaginibacter* OAE612 were significantly diminished (Fig. 7). Aside from strain OAS925, *Rhizobium* OAE497, *Lysobacter* OAE881, and *Rhodococcus* OAS809 were also significantly more enriched in the rhizosphere than in the sand. The results matched well with our observations on the relative abundance (Fig. 6C). These differences between sample types were expected because the *Brachypodium* plant produces compounds that affect soil microbe growth and metabolism (Kawasaki et al. 2016; Sharma et al. 2020). Coker et al. (2022) also observed that *Burkholderia* OAS925 and *Rhizobium* OAE497 increased in relative abundance after growing in the presence of *B. distachyon* (Coker et al. 2022).

Due to the ability of *Variovorax* OAS795 to efficiently outcompete the other SynCom members in the absence of a plant, and the ability of *Burkholderia* OAS925 to efficiently outcompete the rest of SynCom members and competitively colonize plant rhizosphere and roots, these two strains are interesting targets for future research in order to identify the genomic traits that allow such competitive behaviors.

### Inoculation practices affect microbial communities

Inoculation methods are an important factor contributing to plant-microbe interactions, especially when harvest occurs early (Bashan 1998; Kumar et al. 2022; Lopes et al. 2021). However, comprehensive comparisons between different inoculation methods are scarce. The present study demonstrates that inoculation practices significantly shaped the 14 DAG microbial community but not the 21 DPI samples. This points out the importance of implementing different inoculation methods into experimental designs and to the need of comparing multiple inoculation methods when studying plant-microbe interactions. A recent study on sorghum (*Sorghum bicolor*) also showed that inoculation practices impacted the total abundance of different bacterial species at an earlier harvesting point, but not at a later harvesting point (Chai et al. 2022).

Our results from 14 DAG plants suggest that microbial inoculation practices influenced dynamics of plant-microbe interactions, especially in the rhizosphere, before they reached a steady state. Seedling inoculation method (8 DAG) had a significantly higher abundance of *Burkholderia* OAS925 and *Lysobacter* OAE881 and significantly lower *Rhodococcus* OAS809, compared to other inoculation methods at 14 DAG (Fig. 5). This pattern indicates that the seedling inoculation method had the most distinct microbial community compared to the other inoculation methods. The fact that inoculation methods significantly shaped microbial communities may be explained by the dynamic profiles of *B. distachyon* exudate and the response of the SynCom to root exudates (McLaughlin et al. 2023; Saleh et al. 2020; Sharma et al. 2020; Zhalnina et al. 2018). *B. distachyon* exudate profiles can vary significantly within hours (McLaughlin et al. 2023) and weeks (Zhalnina et al. 2018). Thus, the exudate concentration and profile of the 8 DAG group may be significantly different from that of the other groups. It has been shown that *B. distachyon* root exudates affect bacteria in their chemotaxis and gene expression within 48 hours of root exudate addition (Saleh et al. 2020; Sharma et al. 2020). Therefore, the differential abundance of SynCom members observed with the different inoculation methods could be attributed to the SynCom-root exudate interactions.

### Transplant inoculation coupled with a longer co-culture period is the best practice for the EcoFAB-*B. distachyon*-SynCom fabricated ecosystem

This study aimed to establish a protocol that provides the most reproducible and reliable plant-microbe interaction results to serve as a baseline for standardized, small-scale fabricated ecosystem studies. We predict such system would be valuable for multiple downstream experiments including, but not limited to, modeling rhizosphere microbe-microbe and plant-microbe interactions, building a more complex fabricated ecosystem that contains fungi/phages/archaea, and *in-situ* microbial community gene editing. To reach that goal, the system needs to have persistent plant-microbe associations, a reproducible and stable microbial community, non-adversable plant phenotypes, and low labor intensity.

Here, we showed that the SynCom associated with *B. distachyon* persistently for 21 days, much longer than previously reported (Coker et al. 2022). Moreover, the microbial community reached the same steady state at 21 DPI regardless of inoculation method, demonstrating the stability of the fabricated ecosystem established in this study. Among the inoculation methods tested, transplant inoculation (3 DAG group) showed the most reproducible within-group microbial community and a neutral plant growth effect. Transplant inoculation is the most commonly used method for lab studies in part due to its less labor intensity (Herrera Paredes et al. 2018; Hu et al. 2020; Schillaci et al. 2020). We conclude that transplant inoculation coupled with a longer co-culture period (late harvesting point) is a good strategy to conduct bench-top, large-scale fabricated ecosystem studies.

In summary, our results for both plant growth and microbial community abundance suggest that the sand-filled EcoFAB-*B. distachyon*-SynCom system is highly comparable to conventional plant growing methods and thus is a suitable fabricated ecosystem for future studies. Additionally, we showed that transplant inoculation is the best application procedure to study the steady state microbiome in the sand-filled EcoFAB-*B. distachyon*-SynCom system. This research provides a baseline and foundational information for using our fabricated ecosystem to study plant-microbe-environment interactions.

## DATA AVAILABILITY

The16S rRNA gene fragment reads generated through this study can be found on the NCBI BioProject Accession PRJNA975970. All code used in this study can be accessed through Github at https://github.com/JennyMortimer/EcoFAB-Brachypodium-rhizosphere.

## ACKNOWLEDGMENTS

The authors thank John Vogel for providing *Brachypodium distachyon* Bd21-3 seeds and the QB3 Core Research Facility at UC Berkeley for sequencing.

## Literature Cited

Adesemoye, A. O., and Kloepper, J. W. 2009. Plant–microbes interactions in enhanced fertilizer-use efficiency. Appl. Microbiol. Biotechnol. 85:1–12.

Atlas, R. 2010. Handbook of microbiological media, fourth edition. CRC Press.

Bashan, Y. 1998. Inoculants of plant growth-promoting bacteria for use in agriculture. Biotechnol. Adv. 16:729–770.

Bhardwaj, D., Ansari, M. W., Sahoo, R. K., and Tuteja, N. 2014. Biofertilizers function as key player in sustainable agriculture by improving soil fertility, plant tolerance and crop productivity. Microb. Cell Fact. 13:66.

Bragg, J. N., Wu, J., Gordon, S. P., Guttman, M. E., Thilmony, R., Lazo, G. R., Gu, Y. Q., and Vogel, J. P. 2012. Generation and characterization of the Western Regional Research Center *Brachypodium* T-DNA insertional mutant collection. PLoS ONE. 7:e41916.

Busch, J., Mendelssohn, I. A., Lorenzen, B., Brix, H., and Miao, S. 2006. A rhizotron to study root growth under flooded conditions tested with two wetland Cyperaceae. Flora - Morphology, Distribution, Functional Ecology of Plants. 201:429–439.

Chai, Y. N., Futrell, S., and Schachtman, D. P. 2022. Assessment of bacterial inoculant delivery methods for cereal crops. Front. Microbiol. 13:791110.

Coker, J., Zhalnina, K., Marotz, C., Thiruppathy, D., Tjuanta, M., D’Elia, G., Hailu, R., Mahosky, T., Rowan, M., Northen, T. R., and Zengler, K. 2022. A reproducible and tunable synthetic soil microbial community provides new insights into microbial ecology. mSystems. 7:e0095122.

Cole, J. R., Wang, Q., Fish, J. A., Chai, B., McGarrell, D. M., Sun, Y., Brown, C. T., Porras-Alfaro, A., Kuske, C. R., and Tiedje, J. M. 2014. Ribosomal Database Project: data and tools for high throughput rRNA analysis. Nucleic Acids Res. 42:D633–42.

Compant, S., Nowak, J., Coenye, T., Clément, C., and Ait Barka, E. 2008. Diversity and occurrence of *Burkholderia* spp. in the natural environment. FEMS Microbiol. Rev. 32:607– 626.

Donn, S., Kirkegaard, J. A., Perera, G., Richardson, A. E., and Watt, M. 2015. Evolution of bacterial communities in the wheat crop rhizosphere. Environ. Microbiol. 17:610–621.

Draper, J., Mur, L. A., Jenkins, G., Ghosh-Biswas, G. C., Bablak, P., Hasterok, R., and Routledge, A. P. 2001. *Brachypodium distachyon*. A new model system for functional genomics in grasses. Plant Physiol. 127:1539–1555.

Edgar, R. C. 2010. Search and clustering orders of magnitude faster than BLAST. Bioinformatics. 26:2460–2461.

Edgar, R. C. 2016. UNOISE2: improved error-correction for Illumina 16S and ITS amplicon sequencing. BioRxiv.

Finkel, O. M., Salas-González, I., Castrillo, G., Conway, J. M., Law, T. F., Teixeira, P. J. P. L., Wilson, E. D., Fitzpatrick, C. R., Jones, C. D., and Dangl, J. L. 2020. A single bacterial genus maintains root growth in a complex microbiome. Nature. 587:103–108.

Gao, J., Sasse, J., Lewald, K. M., Zhalnina, K., Cornmesser, L. T., Duncombe, T. A., Yoshikuni, Y., Vogel, J. P., Firestone, M. K., and Northen, T. R. 2018. Ecosystem Fabrication (EcoFAB) protocols for the construction of laboratory ecosystems designed to study plant-microbe Interactions. J. Vis. Exp. :e57170.

Hallmann, J., Rodrıguez-Kábana, R., and Kloepper, J. W. 1999. Chitin-mediated changes in bacterial communities of the soil, rhizosphere and within roots of cotton in relation to nematode control. Soil Biol. Biochem. 31:551–560.

Han, J.-I., Choi, H.-K., Lee, S.-W., Orwin, P. M., Kim, J., Laroe, S. L., Kim, T.-G., O’Neil, J., Leadbetter, J. R., Lee, S. Y., Hur, C.-G., Spain, J. C., Ovchinnikova, G., Goodwin, L., and Han, C. 2011. Complete genome sequence of the metabolically versatile plant growth-promoting endophyte *Variovorax paradoxus* S110. J. Bacteriol. 193:1183–1190.

Herrera Paredes, S., Gao, T., Law, T. F., Finkel, O. M., Mucyn, T., Teixeira, P. J. P. L., Salas González, I., Feltcher, M. E., Powers, M. J., Shank, E. A., Jones, C. D., Jojic, V., Dangl, J. L., and Castrillo, G. 2018. Design of synthetic bacterial communities for predictable plant phenotypes. PLoS Biol. 16:e2003962.

Hong, S. Y., Park, J. H., Cho, S. H., Yang, M. S., and Park, C. M. 2011. Phenological growth stages of *Brachypodium distachyon*: codification and description. Weed Res. 51:612–620.

Hsia, M. M., O’Malley, R., Cartwright, A., Nieu, R., Gordon, S. P., Kelly, S., Williams, T. G., Wood, D. F., Zhao, Y., Bragg, J., Jordan, M., Pauly, M., Ecker, J. R., Gu, Y., and Vogel, J. P. 2017. Sequencing and functional validation of the JGI *Brachypodium distachyon* T-DNA collection. Plant J. 91:361–370.

Hu, D., Li, S., Li, Y., Peng, J., Wei, X., Ma, J., Zhang, C., Jia, N., Wang, E., and Wang, Z. 2020. *Streptomyces* sp. strain TOR3209: a rhizosphere bacterium promoting growth of tomato by affecting the rhizosphere microbial community. Sci. Rep. 10:20132.

International Brachypodium Initiative. 2010. Genome sequencing and analysis of the model grass *Brachypodium distachyon*. Nature. 463:763–768.

Kawasaki, A., Donn, S., Ryan, P. R., Mathesius, U., Devilla, R., Jones, A., and Watt, M. 2016. Microbiome and exudates of the root and rhizosphere of *Brachypodium distachyon*, a Model for Wheat. PLoS ONE. 11:e0164533.

Ke, J., Wang, B., and Yoshikuni, Y. 2021. Microbiome engineering: synthetic biology of plant-associated microbiomes in sustainable agriculture. Trends Biotechnol. 39:244–261.

Khatoon, Z., Huang, S., Rafique, M., Fakhar, A., Kamran, M. A., and Santoyo, G. 2020. Unlocking the potential of plant growth-promoting rhizobacteria on soil health and the sustainability of agricultural systems. J. Environ. Manage. 273:111118.

Kumar, S., Diksha, Sindhu, S. S., and Kumar, R. 2022. Biofertilizers: An ecofriendly technology for nutrient recycling and environmental sustainability. Current Research in Microbial Sciences. 3:100094.

Leadbetter, J. R., and Greenberg, E. P. 2000. Metabolism of acyl-homoserine lactone quorum-sensing signals by *Variovorax paradoxus*. J. Bacteriol. 182:6921–6926.

van der Linde, K., Lim, B. T., Rondeel, J. M., Antonissen, L. P., and de Jong, G. M. 1999. Improved bacteriological surveillance of haemodialysis fluids: a comparison between Tryptic soy agar and Reasoner’s 2A media. Nephrol. Dial. Transplant. 14:2433–2437.

Liu, Y.-X., Qin, Y., and Bai, Y. 2019. Reductionist synthetic community approaches in root microbiome research. Curr. Opin. Microbiol. 49:97–102.

Li, X., Rui, J., Mao, Y., Yannarell, A., and Mackie, R. 2014. Dynamics of the bacterial community structure in the rhizosphere of a maize cultivar. Soil Biol. Biochem. 68:392–401.

Lopes, M. J. dos S., Dias-Filho, M. B., and Gurgel, E. S. C. 2021. Successful plant growth-promoting microbes: inoculation methods and abiotic factors. Front. Sustain. Food Syst. 5.

Marín, O., González, B., and Poupin, M. J. 2021. From microbial dynamics to functionality in the rhizosphere: A systematic review of the opportunities with synthetic microbial communities. Front. Plant Sci. 12:650609.

Matos, A., and Garland, J. L. 2005. Effects of community versus single strain inoculants on the biocontrol of Salmonella and microbial community dynamics in alfalfa sprouts. J. Food Prot. 68:40–48.

McCarty, N. S., and Ledesma-Amaro, R. 2019. Synthetic biology tools to engineer microbial communities for biotechnology. Trends Biotechnol. 37:181–197.

McLaughlin, S., Zhalnina, K., Kosina, S., Northen, T. R., and Sasse, J. 2023. The core metabolome and root exudation dynamics of three phylogenetically distinct plant species. Nat. Commun. 14:1649.

Meisner, A., Wepner, B., Kostic, T., van Overbeek, L. S., Bunthof, C. J., de Souza, R. S. C., Olivares, M., Sanz, Y., Lange, L., Fischer, D., Sessitsch, A., Smidt, H., and MicrobiomeSupport Consortium. 2022. Calling for a systems approach in microbiome research and innovation. Curr. Opin. Biotechnol. 73:171–178.

Mitter, E. K., Tosi, M., Obregón, D., Dunfield, K. E., and Germida, J. J. 2021. Rethinking crop nutrition in times of modern microbiology: innovative biofertilizer technologies. Front. Sustain. Food Syst. 5.

Morya, R., Salvachúa, D., and Thakur, I. S. 2020. *Burkholderia*: an untapped but promising bacterial genus for the conversion of aromatic compounds. Trends Biotechnol. 38:963–975.

O’Callaghan, M. 2016. Microbial inoculation of seed for improved crop performance: issues and opportunities. Appl. Microbiol. Biotechnol. 100:5729–5746.

Oburger, E., Dell‘mour, M., Hann, S., Wieshammer, G., Puschenreiter, M., and Wenzel, W. W. 2013. Evaluation of a novel tool for sampling root exudates from soil-grown plants compared to conventional techniques. Environ. Exp. Bot. 87:235–247.

Oleńska, E., Małek, W., Wójcik, M., Swiecicka, I., Thijs, S., and Vangronsveld, J. 2020. Beneficial features of plant growth-promoting rhizobacteria for improving plant growth and health in challenging conditions: A methodical review. Sci. Total Environ. 743:140682.

Pallud, C., Viallard, V., Balandreau, J., Normand, P., and Grundmann, G. 2001. Combined use of a specific probe and PCAT medium to study *Burkholderia* in soil. J. Microbiol. Methods. 47:25–34.

Parada, A. E., Needham, D. M., and Fuhrman, J. A. 2016. Every base matters: assessing small subunit rRNA primers for marine microbiomes with mock communities, time series and global field samples. Environ. Microbiol. 18:1403–1414.

Parke, J. L., and Gurian-Sherman, D. 2001. Diversity of the *Burkholderia cepacia* complex and implications for risk assessment of biological control strains. Annu. Rev. Phytopathol. 39:225–258.

Parnell, J. J., Berka, R., Young, H. A., Sturino, J. M., Kang, Y., Barnhart, D. M., and DiLeo, M. V. 2016. From the lab to the farm: an industrial perspective of plant beneficial microorganisms. Front. Plant Sci. 7:1110.

Pradhan, S., Tyagi, R., and Sharma, S. 2022. Combating biotic stresses in plants by synthetic microbial communities: Principles, applications and challenges. J. Appl. Microbiol. 133:2742–2759.

Price, M. N., Wetmore, K. M., Waters, R. J., Callaghan, M., Ray, J., Liu, H., Kuehl, J. V., Melnyk, R. A., Lamson, J. S., Suh, Y., Carlson, H. K., Esquivel, Z., Sadeeshkumar, H., Chakraborty, R., Zane, G. M., Rubin, B. E., Wall, J. D., Visel, A., Bristow, J., Blow, M. J., and Deutschbauer, A. M. 2018. Mutant phenotypes for thousands of bacterial genes of unknown function. Nature. 557:503–509.

Quince, C., Lanzen, A., Davenport, R. J., and Turnbaugh, P. J. 2011. Removing noise from pyrosequenced amplicons. BMC Bioinformatics. 12:38.

Reasoner, D. J., and Geldreich, E. E. 1985. A new medium for the enumeration and subculture of bacteria from potable water. Appl. Environ. Microbiol. 49:1–7.

Rocha, I., Ma, Y., Souza-Alonso, P., Vosátka, M., Freitas, H., and Oliveira, R. S. 2019. Seed coating: A tool for delivering beneficial microbes to agricultural crops. Front. Plant Sci. 10:1357.

Saleh, D., Sharma, M., Seguin, P., and Jabaji, S. 2020. Organic acids and root exudates of *Brachypodium distachyon*: effects on chemotaxis and biofilm formation of endophytic bacteria. Can. J. Microbiol. 66:562–575.

dos Santos, S. G., Chaves, V. A., da Silva Ribeiro, F., Alves, G. C., and Reis, V. M. 2019. Rooting and growth of pre-germinated sugarcane seedlings inoculated with diazotrophic bacteria. Appl. Soil Ecol. 133:12–23.

Sarkar, D., Singh, S., Parihar, M., and Rakshit, A. 2021. Seed bio-priming with microbial inoculants: A tailored approach towards improved crop performance, nutritional security, and agricultural sustainability for smallholder farmers. CRSUST. 3:100093.

Sasse, J., Kant, J., Cole, B. J., Klein, A. P., Arsova, B., Schlaepfer, P., Gao, J., Lewald, K., Zhalnina, K., Kosina, S., Bowen, B. P., Treen, D., Vogel, J., Visel, A., Watt, M., Dangl, J. L., and Northen, T. R. 2019. Multilab EcoFAB study shows highly reproducible physiology and depletion of soil metabolites by a model grass. New Phytol. 222:1149–1160.

Sasse, J., Kosina, S. M., de Raad, M., Jordan, J. S., Whiting, K., Zhalnina, K., and Northen, T. R. 2020. Root morphology and exudate availability are shaped by particle size and chemistry in *Brachypodium distachyon*. Plant Direct. 4:e00207.

Schillaci, M., Arsova, B., Walker, R., Smith, P. M. C., Nagel, K. A., Roessner, U., and Watt, M. 2020. Time-resolution of the shoot and root growth of the model cereal *Brachypodium* in response to inoculation with *Azospirillum* bacteria at low phosphorus and temperature. Plant Growth Regul.

Schlemper, T. R., Leite, M. F. A., Lucheta, A. R., Shimels, M., Bouwmeester, H. J., van Veen, J. A., and Kuramae, E. E. 2017. Rhizobacterial community structure differences among sorghum cultivars in different growth stages and soils. FEMS Microbiol. Ecol. 93.

Sharma, M., Saleh, D., Charron, J.-B., and Jabaji, S. 2020. A crosstalk between *Brachypodium* root exudates, organic acids, and *Bacillus velezensis* B26, a growth promoting bacterium. Front. Microbiol. 11:575578.

Sharpless, W., Sander, K., Song, F., Kuehl, J., and Arkin, A. P. 2022. Towards environmental control of microbiomes. BioRxiv.

Shayanthan, A., Ordoñez, P. A. C., and Oresnik, I. J. 2022. The role of synthetic microbial communities (syncom) in sustainable agriculture. Front. Agron. 4.

Song, F., Kuehl, J. V., Chandran, A., and Arkin, A. P. 2021. A simple, cost-effective, and automation-friendly direct PCR approach for bacterial community analysis. mSystems. 6:e0022421.

de Souza, R. S. C., Armanhi, J. S. L., and Arruda, P. 2020. From microbiome to traits: designing synthetic microbial communities for improved crop resiliency. Front. Plant Sci. 11:1179.

Tyagi, R., Pradhan, S., Bhattacharjee, A., Dubey, S., and Sharma, S. 2022. Management of abiotic stresses by microbiome-based engineering of the rhizosphere. J. Appl. Microbiol.

United Nation. 2022. The Sustainable Development Goals Report 2022. ed. Lois Jensen. United Nations.

Valetti, L., Iriarte, L., and Fabra, A. 2018. Growth promotion of rapeseed (*Brassica napus*) associated with the inoculation of phosphate solubilizing bacteria. Appl. Soil Ecol. 132:1– 10.

Vogel, J., and Hill, T. 2008. High-efficiency *Agrobacterium*-mediated transformation of *Brachypodium distachyon* inbred line Bd21-3. Plant Cell Rep. 27:471–478.

Vogel, J. P., ed. 2016. Genetics and genomics of Brachypodium. Cham: Springer International Publishing.

Watt, M., Schneebeli, K., Dong, P., and Wilson, I. W. 2009. The shoot and root growth of *Brachypodium* and its potential as a model for wheat and other cereal crops. Functional Plant Biol. 36:960.

Yee, M. O., Kim, P., Li, Y., Singh, A. K., Northen, T. R., and Chakraborty, R. 2021. Specialized plant growth chamber designs to study complex rhizosphere interactions. Front. Microbiol. 12:625752.

York, L. M., Cumming, J. R., Trusiak, A., Bonito, G., Haden, A. C., Kalluri, U. C., Tiemann, L. K., Andeer, P. F., Blanc-Betes, E., Diab, J. H., Favela, A., Germon, A., Gomez-Casanovas, N., Hyde, C. A., Kent, A. D., Ko, D. K., Lamb, A., Missaoui, A. M., Northen, T. R., Pu, Y., and Yang, W. H. 2022. Bioenergy Underground: Challenges and opportunities for phenotyping roots and the microbiome for sustainable bioenergy crop production. Plant phenome j. 5:e20028.

Zengler, K., Hofmockel, K., Baliga, N. S., Behie, S. W., Bernstein, H. C., Brown, J. B., Dinneny, J. R., Floge, S. A., Forry, S. P., Hess, M., Jackson, S. A., Jansson, C., Lindemann, S. R., Pett-Ridge, J., Maranas, C., Venturelli, O. S., Wallenstein, M. D., Shank, E. A., and Northen, T. R. 2019. EcoFABs: advancing microbiome science through standardized fabricated ecosystems. Nat. Methods. 16:567–571.

Zhalnina, K., Louie, K. B., Hao, Z., Mansoori, N., da Rocha, U. N., Shi, S., Cho, H., Karaoz, U., Loqué, D., Bowen, B. P., Firestone, M. K., Northen, T. R., and Brodie, E. L. 2018. Dynamic root exudate chemistry and microbial substrate preferences drive patterns in rhizosphere microbial community assembly. Nat. Microbiol. 3:470–480.

Zhang, J., Kobert, K., Flouri, T., and Stamatakis, A. 2014. PEAR: a fast and accurate Illumina Paired-End reAd mergeR. Bioinformatics. 30:614–620.

