## Supplemental figures and tables for "Impact of inoculation practices on microbiota assembly and community stability in a fabricated ecosystem"

### Supplemental Tables

**Supplemental Table S1. *B. distachyon* developmental stage in EcoFABs filled with hydroponic media with published data.** The *B. distachyon* developmental stage was recorded at different days after germination (DAG) with a brief description. The description is compared with the one published by (Hong et al. 2011) that used the BBCH (Biologische Bundesanstalt, Bundessortenamt und CHemische Industrie) numerical scale system (Bleiholder et al. 2001).

| Days after germination (DAG) | Description | BBCH scale | Days in Hong et al. 2011 |
|---|---|---|---|
| 0 | Germination | Stage 0 | 0 |
| 3 | Coleoptile emerged | Stage 07 | 2.9 |
| 8 | First leaf unfolded | Stage 11 | 7.6 |
| 14 | Beginning of tillering | Stage 21 | 15.8 |
| 28 | Flag leaf sheath extending | Stage 41 | 24.9 |

**Supplemental Table S2. Permutational multivariate analysis of variance (PERMANOVA) results of Bray-Curtis dissimilarity for all data collected at 14 days after germination (DAG).** Samples separated primarily by sample type (Sample_Type) and moderately by inoculation method (Inoc_Day), both combined to explain 45% of the variation. When separated by sample type, inoculation practices significantly affected the microbial community in the rhizosphere, moderately in sand, but not in the root. Significance codes of the P: 0 '***' 0.001 '**' 0.01 '*' 0.05 '.' 0.1 ' ' 1.

| Factor | | Degrees of Freedom | Sum of Squares | R square | F value | P value | Significant codes |
|---|---|---|---|---|---|---|---|
| All Data | Sample_Type | 2 | 6.0452 | 0.4227 | 21.9244 | 0.0001 | *** |
| | Inoc_Day | 2 | 0.5299 | 0.0371 | 1.9218 | 0.0585 | . |
| | Sample_Type: Inoc_Day | 4 | 0.5586 | 0.0391 | 1.0129 | 0.4283 | |
| | Residual | 52 | 7.1690 | 0.5012 | | | |
| | Total | 60 | 14.3027 | 1.0000 | | | |
| Sand | Inoc_Day | 2 | 0.3140 | 0.1896 | 1.9892 | 0.0831 | . |
| | Residual | 17 | 1.3415 | 0.8104 | | | |
| | Total | 19 | 1.6555 | 1.0000 | | | |
| Rhizo-sphere | Inoc_Day | 2 | 0.4527 | 0.2786 | 3.2824 | 0.0060 | ** |

| | | | | | | |
|---|---|---|---|---|---|---|
| | Residual | 17 | 1.1724 | 0.7214 | | |
| | Total | 19 | 1.6251 | 1.0000 | | |
| Root | Inoc_Day | 2 | 0.3218 | 0.0647 | 0.6222 | 0.7154 |
| | Residual | 18 | 4.6551 | 0.9353 | | |
| | Total | 20 | 4.9769 | 1.0000 | | |

**Supplemental Table S3. Permutational multivariate analysis of variance (PERMANOVA) results of Bray-Curtis dissimilarity for all data collected at 21 days post-inoculation (DPI).** Samples separated primarily by sample type (Sample_Type) but not by inoculation method (Inoc_Day). Significance codes of the P: 0 '***' 0.001 '**' 0.01 '*' 0.05 '.' 0.1 ' ' 1.

| Factor | Degrees of Freedom | Sum of Squares | R square | F value | P value | Significant codes |
|---|---|---|---|---|---|---|
| Sample_Type | 2 | 6.325 | 0.538 | 32.163 | 0.000 | *** |
| Inoc_Day | 2 | 0.304 | 0.026 | 1.544 | 0.147 | |
| Sample_Type:Inoc_Day | 4 | 0.501 | 0.043 | 1.274 | 0.211 | |
| Residual | 47 | 4.621 | 0.393 | | | |
| Total | 55 | 11.751 | 1.000 | | | |

**Supplemental figures**

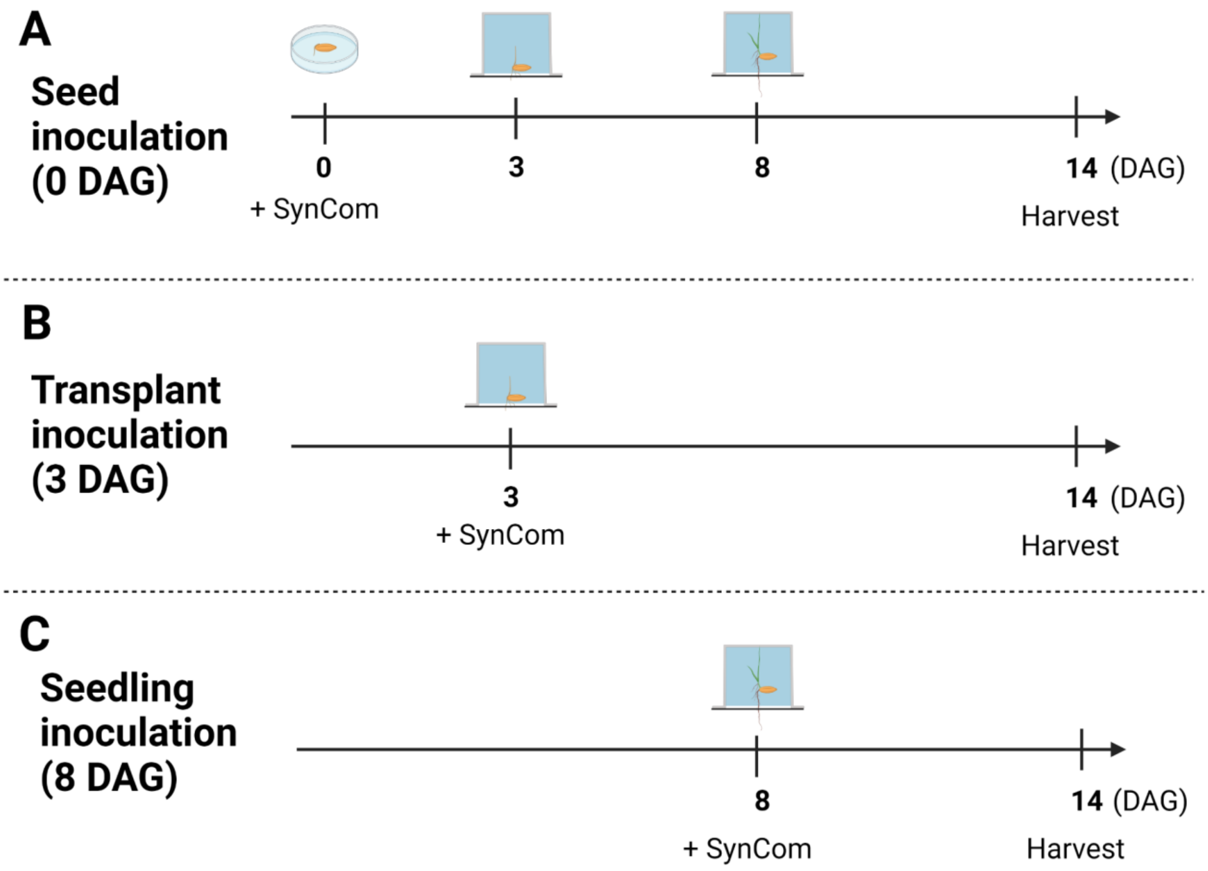


**Supplemental Fig. S1.** Overview of different SynCom inoculation practices on *B. distachyon* plants and harvesting when the plant reached 14 days after germination (DAG). The method of application of the SynCom to the plants was **A**, seed inoculation (0 DAG), **B**, transplant inoculation (3 DAG), **C**, seedling inoculation (8 DAG).

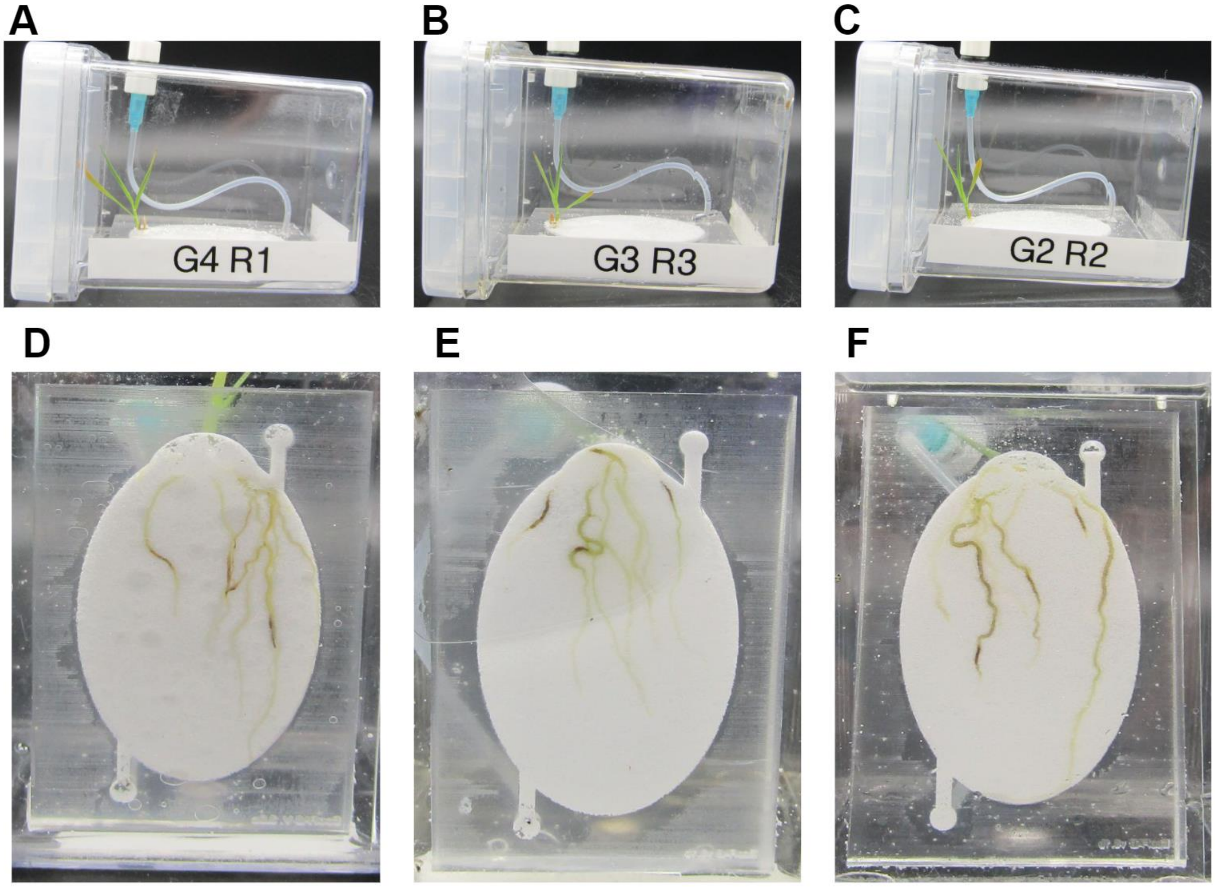


**Supplemental Fig. S2.** Fourteen days after germination (DAG) *B. distachyon* plants grown in sand-filled EcoFABs inoculated with the SynCom using different methods. **A**, **D**, 0 DAG (seed inoculation); **B**, **E**, 3 DAG (transplant inoculation); **C**, **F**, 8 DAG (seedling inoculation). Plants were harvested at 14 DAG.

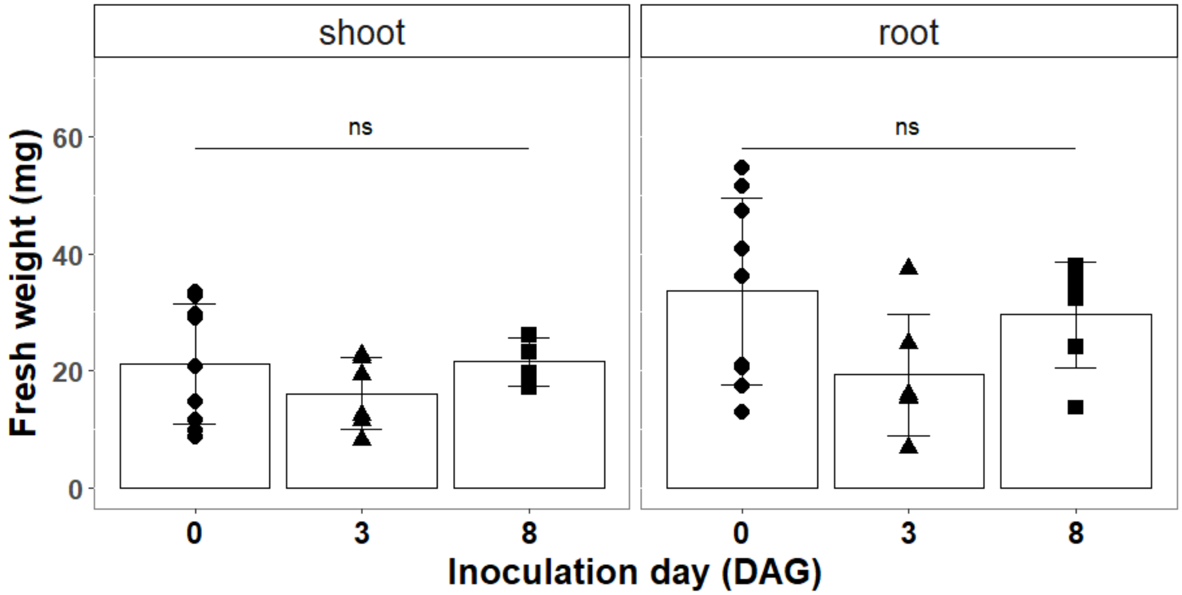


**Supplemental Fig. S3.** *B. distachyon* plant fresh weight 14 days after germination (DAG). Shoot and root weight was similar between different inoculation practices. ANOVA *P* value was 0.406 for the shoot weight and 0.134 for the root weight. Biological replicates were six or more in each group.

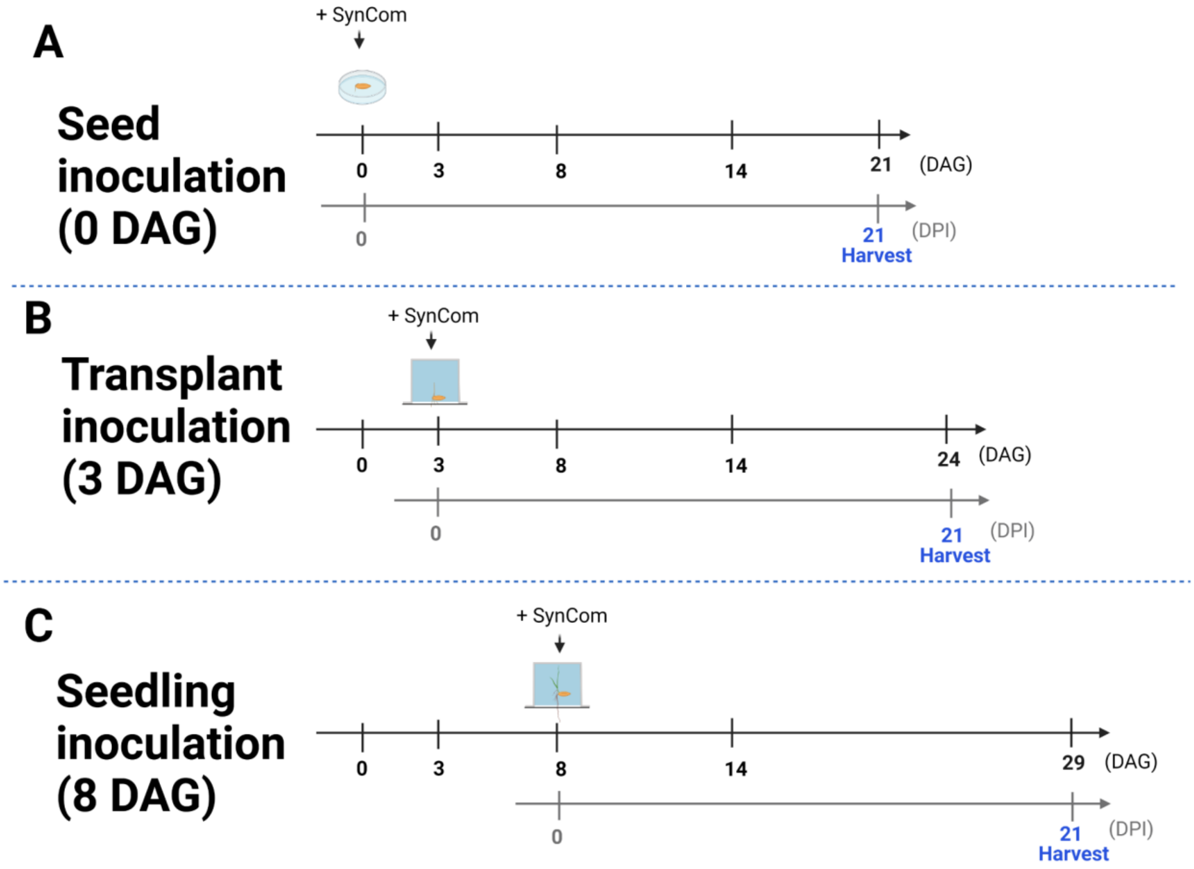


**Supplemental Fig. S4.** Overview of different SynCom inoculation practices on *B. distachyon* plants and harvesting 21 days post-inoculation (DPI). The method of application of the SynCom to the plant was **A**, seed inoculation (0 days after germination; DAG); **B**, transplant inoculation (3 DAG); **C**, seedling inoculation (8 DAG).

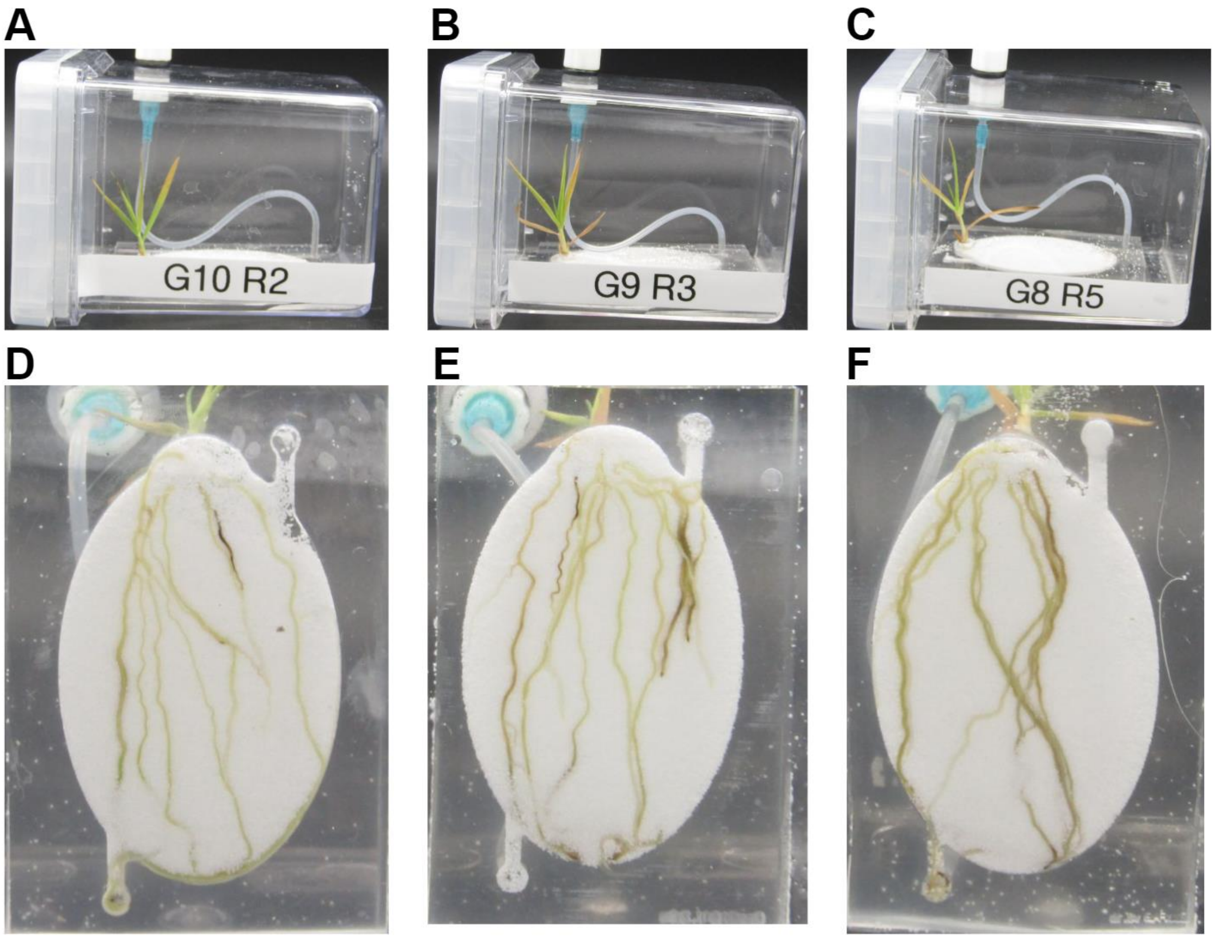


**Supplemental Fig. S5.** *B. distachyon* grown in sand-filled EcoFABs inoculated with the SynCom using different methods. **A**, **D**, 0 days after germination (DAG; seed inoculation); **B**, **E**, 3 DAG (transplant inoculation); **C**, **F**, 8 DAG (seedling inoculation). Plants were harvested 21 days post-inoculation (DPI).

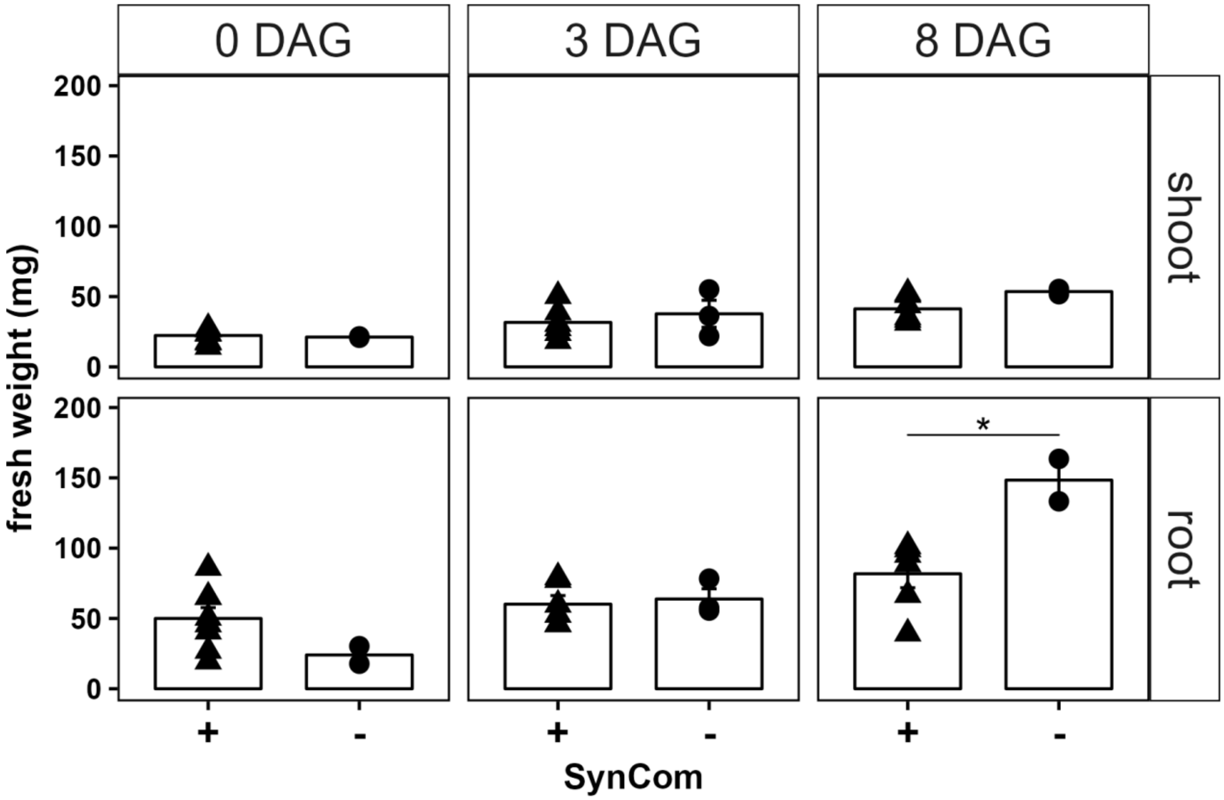


**Supplemental Fig. S6.** *B. distachyon* plant fresh weight 21 days post-inoculation (DPI). SynCom inoculation was performed when *B. distachyon* was at 0 days after germination (DAG), 3 DAG, 8 DAG. Three non-inoculated *B. distachyon* were also grown in parallel as controls (denoted as “-”). SynCom inoculation significantly lowered the root biomass in the 8 DAG groups but not in shoot biomass. On the other hand, the plant biomass in the 0 DAG and 3 DAG groups were not significantly affected by SynCom inoculation compared to the control. Statistical analyses involved a t-test (alpha = 0.05) between the SynCom inoculated (+) and non-inoculated (-) plants of each group. Significance codes of the P: 0 ‘***’ 0.001 ‘**’ 0.01 ‘*’ 0.05 ‘.’ 0.1 ‘ ’ 1.
